# A neuronal pathway for the supraspinal relay of warm and cool sensation

**DOI:** 10.64898/2026.01.19.700348

**Authors:** Xinying Zhang, Farin Bourojeni, Samuel Ferland, Anna McFarlane, Kieran A. Boyle, Madoka Koyanagi, Marie-Eve Paquet, Andrew J. Todd, Junichi Hachisuka, Yves De Koninck, Feng Wang, Artur Kania

## Abstract

Cutaneous thermosensation is a fundamental sense allowing discrimination of external temperature changes and driving adaptive responses essential for survival and well-being. This sensory process begins with the activation of peripheral thermoreceptors, which encode the quality and intensity of thermal stimuli into neuronal activity that is further integrated by spinal cord circuits and relayed to supraspinal centres through the anterolateral tract (ALT). While classical experiments identified thermal responsive ALT neurons in the superficial spinal dorsal horn, the functional logic of innocuous temperature relay remains poorly understood. We show that, in mice, the majority of Phox2a-lineage lamina I ALT are selectively tuned to innocuous warming or cooling, and receive direct inputs from peripheral thermoreceptors. Their genetic ablation, chemogenetic or optogenetic manipulation impaired temperature discrimination. Our findings contrast with the previous finding that the majority of ALT neurons being mechano- and thermo-nociceptive and provide novel evidence for the existence of a large ensemble of genetically defined warming- and cooling- responsive ALT neurons. The large representation of innocuous temperature in supraspinal-relaying ALT neurons indicates substantial spinal amplification of thermal information and underscores the importance of innocuous thermosensation in the ascending sensory pathway.

## Main

Mammalian spinal cord ALT neurons serve as a hub for the relay of somatosensory information, including mechanosensation, thermosensation, and nociception^1^. ALT neurons within the lamina I of the spinal dorsal horn respond to a variety of somatosensory stimuli, and classical physiological and morphological studies as well as recent *in vivo* calcium imaging experiments argue for the existence of at least cold-selective, nociceptive, and polymodal subtypes^2–6^. While some thermosensory lamina I ALT neurons populations have been identified based on their response to noxious cold and hot stimuli, the integration and relay of innocuous temperatures, which are encountered far more frequently than hot or cold extremes, remains poorly understood^7–10^. In mice, approximately half of ALT neurons are defined by the embryonic expression of the transcription factor Phox2a. Previous studies demonstrated that a subset of these show increased phosphorylation of ERK (pERK) in response to noxious heat, and are located near the termini of Trpm8 and Trpv1 thermoreceptor afferents in lamina I, suggesting a thermosensory function^11,12^.

### Genetically defined thermosensory ALT neurons

We first examined whether mouse Phox2a lamina I (Phox2a^lamI^) ALT neurons constitute a distinct subset of all lamina I ALT neurons by assessing their expression of molecular markers. To visualize mouse Phox2a^lamI^ neurons, we produced Phox2a^tdTomato^ mice by crossing *Phox2a::Cre*^11^ mice with Cre-dependent tdTomato reporter (*R26:lsl-tdTomato*) mice. To distinguish between Phox2a ALT neurons and non-Phox2a ALT neurons, we took advantage of the fact that essentially all ALT neurons innervate the lateral parabrachial nucleus (lPBN) in rodents^13^. Thus, in *R26:lsl-tdTomato*; *Phox2a::Cre* mice bilaterally injected with retrograde tracer, cholera toxin subunit B (CTB), in the lPBN, Phox2a^lamI^ neurons were co-labelled by tdTomato and CTB, whereas non-Phox2a^lamI^ ALT neurons were only labelled by CTB (Fig. 1a, Extended Data Fig. 1a). First, we examined the expression of the transcription factor Zfhx3, proposed to label all long-range spinal projection neurons, including ALT neurons^14^. As expected, essentially all CTB+ lamina I ALT neurons expressed Zfhx3 (Fig. 1c, f). Next, we investigated protein and mRNA markers of ALT neurons that correlate with their proposed somatosensory functions. NK1R, the receptor of substance P, and Tac1, a gene coding for the precursor protein of substance P, have been the predominant marker of ALT neurons and associated with nociception^15^. Aligning with observations in adult and embryonic mice^11,16^, we observed lower levels of NK1R protein, and lower *Tac1* mRNA levels in Phox2a^lamI^ neurons, compared to non-Phox2a^lamI^ ALT neurons (Fig. 1b-f, Extended Data Fig. 1d, e and Supplementary Table 1). Past studies also suggested that ALT neurons can be subdivided by their expression of Calbindin protein and *Tacr1* and *Gpr83* mRNA^17,18^. Of these, we found only *Gpr83* showed preferential expression in Phox2a^lamI^ *versus* non-Phox2a^lamI^ ALT neurons (Extended Data Fig. 1b, c, e and Supplementary Table 1). Since low or no NK1R expression in lamina I ALT neurons has been associated with tuning to cold stimuli^19^, and Tac1 expression with mechanical nociception^20–22^, we hypothesised that Phox2a lamina I ALT neurons may have a preferential role in thermosensation given their low expression of both NK1R and *Tac1*.

**Fig. 1.**
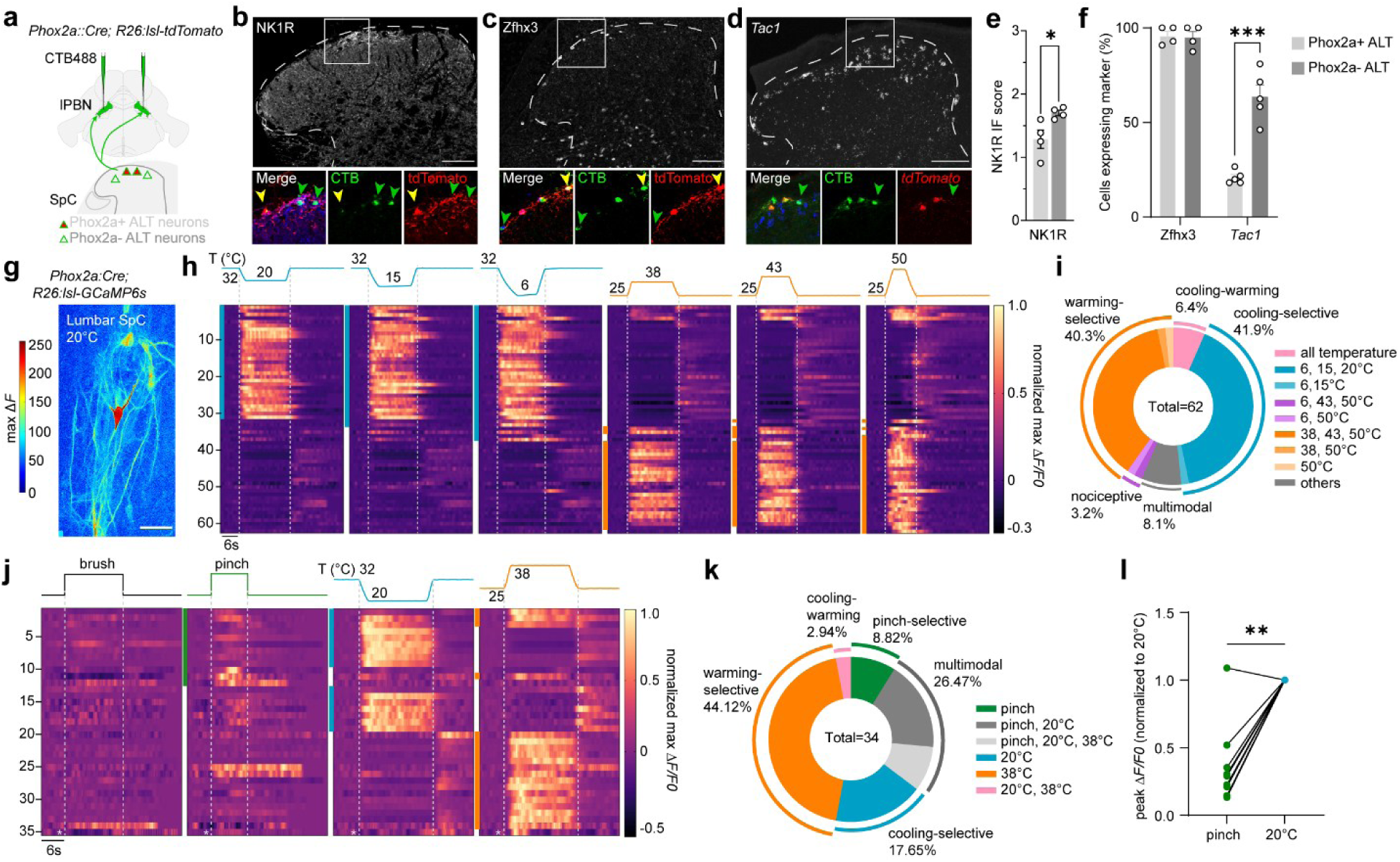
Spinal Phox2a^lamI^ ALT neurons form ascending sensory pathway preferentially tuned to temperature. **a,** Retrograde labelling of ALT neurons by injection of CTB in the bilateral lPBN in *Phox2a::Cre; R26:lsl-tdTomato* mice. **b-d,** Representative images of Phox2a+ (yellow arrowhead) and Phox2a- (green arrowhead) ALT neurons in lumbar spinal cord, labelled with CTB (green) and protein expression of tdTomato (red), and NK1R **(b)** or Zfhx3 (white, **c**) or mRNA expression of *tdTomato* (red) and *Tac1* (white, **d**). **e, f**, Quantification of NK1R expression level **(e)** and fraction of Phox2a+ ALT neurons (light grey) or Phox2a- ALT neurons (dark grey) co-expressing Zfhx3 or Tac1 **(f)**. **g,** Representative imaging field, pseudo coloured based on maximum ΔF of calcium responses to 20°C thermal stimuli in a *Phox2a::Cre; R26:lsl-GCaMP6s* mouse. **h,** Heatmaps of normalized response amplitudes (ΔF/F) to thermal stimuli delivered by a Peltier device: cooling stimuli with baseline temperature of 32°C and decreasing to 20°C, 15°C, or 6°C; warming stimuli with baseline temperature of 25°C and increasing to 38°C, 43°C, or 50°C. Each row represents responses of the same individual neuron (n=62). **i,** Response profile to all thermal stimuli of Phox2a^lamI^ neurons in **(h)**. **j,** Heatmaps of normalized response amplitudes (ΔF/F) to manual brush, pinch, or innocuous thermal stimuli. Each row of the heatmap represents responses of an individual neuron (n=35). **k,** Summary of responses of Phox2a^lamI^ neurons to all stimuli modalities from **(j)**. **l,** Peak calcium signals to pinch of all pinch and 20°C responsive neurons compared to their normalized signals to 20°C. Scale bar: 100 μm in **(b-d) and** 50 μm in **(g)**. Statistics: values plotted as mean ± SEM. N=4 for Zfhx3 and NK1R, N=5 for Tac1, Phox2a+ and Phox2a- ALT neurons. Mean ± SEM. *p <0.05, **p <0.01, ***p < 0.001, non-significant not labelled. Student t-test **(e, f)**, and Paired t-test **(l)**.

To test this idea, we performed functional 2-photon calcium imaging of lumbar Phox2a^lamI^ neurons expressing the GCaMP6s calcium sensor in anesthetized mice (*Phox2a::Cre*; *R26:lsl-GCaMP6s*). A feedback-controlled probe was used to stimulate the plantar side of the hindpaw with a series of fast-ramp-and-hold thermal stimuli, covering innocuous cool (20°C and 15°C) to noxious cold (6°C), as well as innocuous warmth (38°C) to noxious heat (43°C and 50°C) temperature ranges, from adaptive temperature at 32°C or 25°C respectively (Fig. 1g, h, Extended Data Fig. 2a)^23^. We identified a total of 62 Phox2a^lamI^ neurons (ten imaging fields in eight animals) that showed robust Ca^2+^ responses to either cooling or warming, or both. Of these, 41.9% responded to cooling only (cooling-selective), 40.3% responded to warming only (warming-selective), and 6.5% responded to all cooling and warming testing temperatures (all-temp-sensitive, Fig. 1i). These observations provide first direct evidence that Phox2a^lamI^ ALT neurons are indeed thermosensory and comprise two largely complementary subsets responding to innocuous cooling or warming.

Next, we examined whether Phox2a^lamI^ neurons are preferentially tuned to thermal stimuli by comparing their responses to innocuous cooling and warming versus mechanical stimuli (Fig. 1j). Repetitive innocuous brushing and noxious pinching were applied manually to the paw. Of 35 Phox2a^lamI^ neurons tested (six fields from five animals), none responded to brush, 12 responded to pinch (on average 35.0% per animal), 16 responded to cooling (on average 41.6% per animal), and 19 responded to warming (on average 57.5% per animal), while one responded to none of the four tested stimuli (Fig. 1j, Extended Data Fig. 2c, d). Of the 34 responsive neurons, around half thermal sensitive ones responded to cooling or warming (Fig. 1k), consisting with our observation above (Fig. 1i). Specifically, twenty-four neurons were modality-specific, including six neurons responding only to cooling (cooling-selective), 16 neurons responding only to warming (warming-selective), and only three neurons responding only to pinch (pinch-selective, Fig. 1k). ten neurons were multimodal, where one responded to both cooling and warming, and nine responded to both mechanical and thermal stimuli. Notably, all multimodal pinch-responsive neurons also responded to cooling, and they exhibited much smaller Ca^2+^ responses to pinch comparing to their responses to cooling, suggesting that they are still preferentially tuned to thermal stimuli (Fig. 1l). Together, our results suggest that Phox2a lamina I ALT neurons are preferentially tuned to innocuous cool and warm, but not mechanical stimuli.

### Cool and warm encoding by Phox2a ALT neurons

We next considered three potential neuronal coding mechanisms of Phox2a^lamI^ neurons across the innocuous and noxious temperature ranges: (1) tuning to specific temperatures and selective recruitment; (2) recruitment of increasing numbers of neurons at more extreme temperatures; or (3) graded response amplitudes to different temperatures (Fig. 2a)^24^. To distinguish between these, we measured the proportion of thermal-responsive Phox2a^lamI^ neurons that were recruited at various temperatures, which were similar within the innocuous and noxious range (Fig. 2b). In addition, of the cooling-selective neurons, most neurons that responded to a specific cooling temperature also responded to other cooling temperatures, and the same was true for warming-selective neurons (Fig. 2c), together suggesting broad and not specific tuning to both cooling and warming in Phox2a^lamI^ neurons. Furthermore, the peak Ca^2+^ changes of Phox2a^lamI^ neurons did not show a graded increase between innocuous and noxious ranges (Fig. 2d), suggesting that these neurons are broadly tuned to innocuous temperatures, and do not encode temperature changes in the noxious range.

**Fig. 2.**
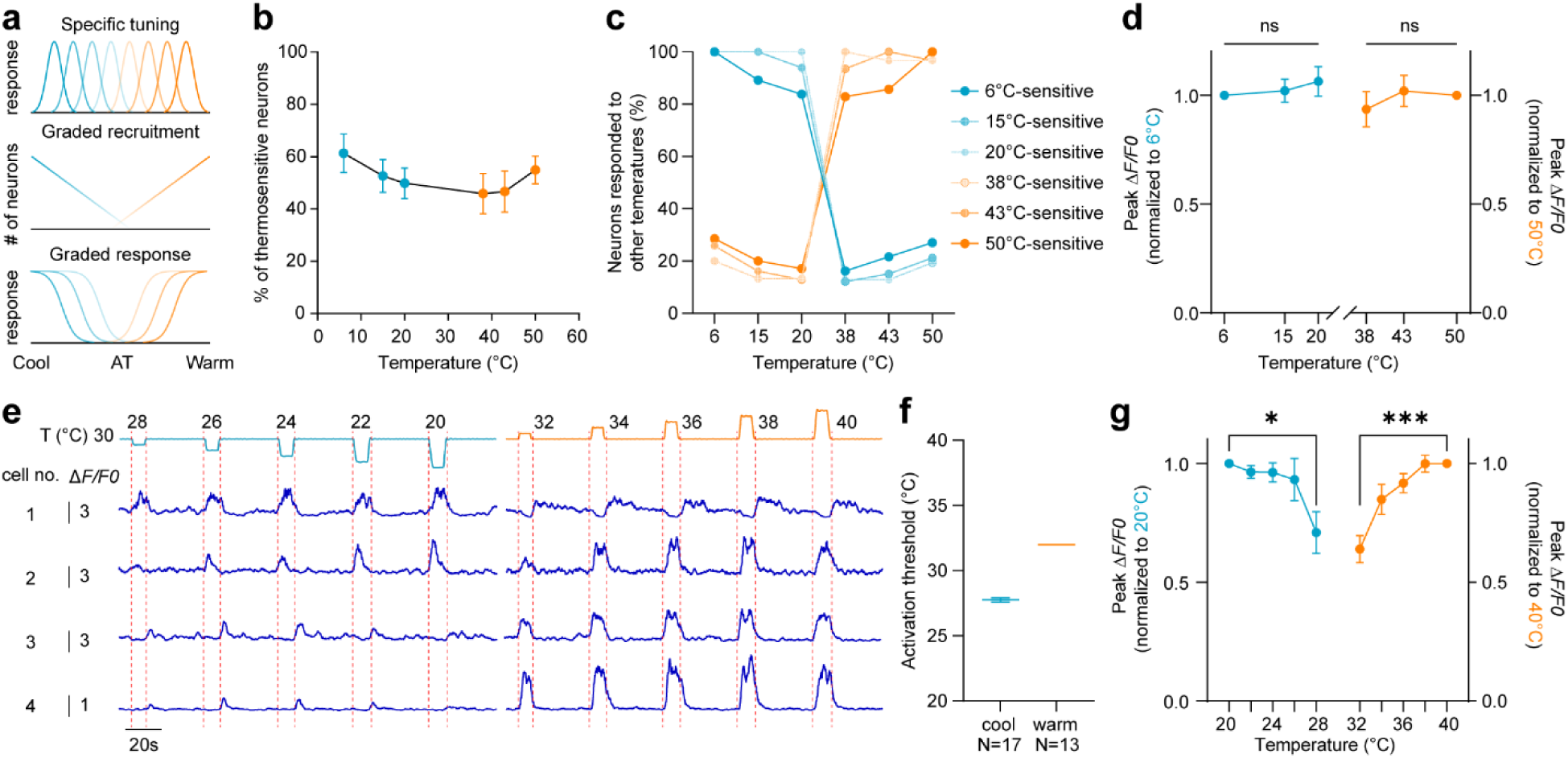
Encoding of cool and warm by Phox2a^lamI^ ALT neurons. **a,** Three possible temperature encoding models adapted from Vestergaard, et al. ^24^ **b, c,** Percentage of all thermosensory neurons that responded to each thermal stimuli **(b)** and percentage of neurons activated by a given temperature (coloured lines) that responded to each thermal stimuli **(c)** showed broad tuning property. There was no significant difference among the responses to different stimuli. **d,** Thermal-sensitive Phox2a^LamI^ neurons do not exhibit stronger response to noxious thermal stimuli compared to innocuous thermal stimuli. **e,** Representative calcium curves from four neurons showing typical responses to step changes in thermal stimuli with a baseline temperature of 30°C. Cell number 1 responded to warming stimuli and to the return phase of coolinging stimuli. Cell number 2 responded to both cooling and warming stimuli. Cell number 3 and 4 responded to warming stimuli and to the return phase from cooling stimuli. **f,** Activation threshold for the thermal induced calcium responses in cooling or warming responsive cells. **g,** Normalized peak amplitude of wide-range cooling neurons (blue, N=17) and wide-range warming neurons (orange, N=13) to decreasing and increasing temperature, respectively. Mean ± SEM. *p <0.05, **p <0.01, ***p < 0.001, ns: non-significant. Ordinary one-way ANOVA **(c)**, Repeated Measurement one-way ANOVA **(d, g)**.

To elucidate the coding strategy of Phox2a^lamI^ neurons in the innocuous temperature range, we examined their Ca^2+^ responses in a series of step-increments or decrements from the adaptive temperature of 30°C (Fig. 2e). Notably, of 27 neurons examined in total, most responded to small increment or decrement of 2°C. Particularly, 16 cooling-responsive neurons were activated at 28°C and one at 26°C (17 neurons in total), and all warming-responsive neurons were activated at 32°C (13 neurons in total), indicating that Phox2a^lamI^ neurons are highly responsive to temperature changes (Fig. 2f). Moreover, both cooling- and warming-responsive neurons showed graded responses within the innocuous range, and their peak Ca^2+^ changes plateaued within the innocuous range, further arguing for a limited ability of these neurons to encode noxious temperatures (Fig. 2d, g). Together, these results demonstrate that Phox2a lamina I ALT neurons are broadly tuned to wide-range cooling or warming, and encode innocuous, but not noxious, temperature changes with graded responses.

### Thermoreceptor inputs to Phox2a neurons

Cutaneous somatosensory information is transmitted to the spinal cord by primary sensory neurons tuned to one or more specific modalities. We reasoned that thermosensory Phox2a ALT neurons may receive direct inputs from thermoreceptors since a previous study reported that Trpm8 cool-thermoreceptor afferents contact the dendrites of Phox2a^lamI^ neurons^25,26^. Additionally, most thermosensory Phox2a^lamI^ neurons were slow-adapting (Fig. 1h), with their normalized full-width-half-maximum (FWHM) of the calcium response larger than 0.5, echoing the characteristics of slow-adapting (SA) Trpm8+ thermoreceptors (Extended Data Fig. 2b; Trpm8 data not shown)^27^. To map direct sensory afferent inputs to Phox2a ALT neurons, we performed monosynaptic retrograde tracing, using an EnvA-pseudotyped, glycoprotein-deleted rabies virus (*EnvA(dG)-RabV::mCherry*) injected unilaterally into the lumbar enlargement of mice expressing the EnvA receptor (TVA) in Phox2a ALT neurons (*Phox2a::Cre; hCdx2:FlpO; R26:lsl-fsf-HTB*, Fig. 3a). As expected, the rabies virus selectively infected spinal dorsal horn Phox2a ALT neurons (mCherry+GFP+, starter cells), but not neurons lacking TVA (Fig. 3b, Extended Data Fig 3a, b). Across L3-L5 dorsal root ganglia (DRGs), we observed rabies-infected sensory neurons (mCherry+ and NeuN+), encompassing peptidergic C- and Aδ-fiber populations (TrkA+, CGRP+) known to include thermoreceptors and nociceptors presynaptic to superficial ALT neurons (Fig. 3c, d, Extended Data Fig. 3c, d)^1^. Among these, we identified neurons expressing *Trpm8 and Trpv1* mRNAs, markers of cold and hot thermoreceptors, respectively, suggesting direct monosynaptic input from both to Phox2a ALT neurons. Given that Trpm8+ and Trpv1+ afferents terminate predominantly in the superficial dorsal horn^25,26,28^, it is likely that these labelling by rabies transfer arise superficially from Phox2a ALT neurons in lamina I rather than deeper layers. In contrast, no non-peptidergic sensory neurons (IB4+) were labelled. However, this result should be interpretated with caution, as this population has well reported to be resistant to rabies infection^29^.

**Figure 3.**
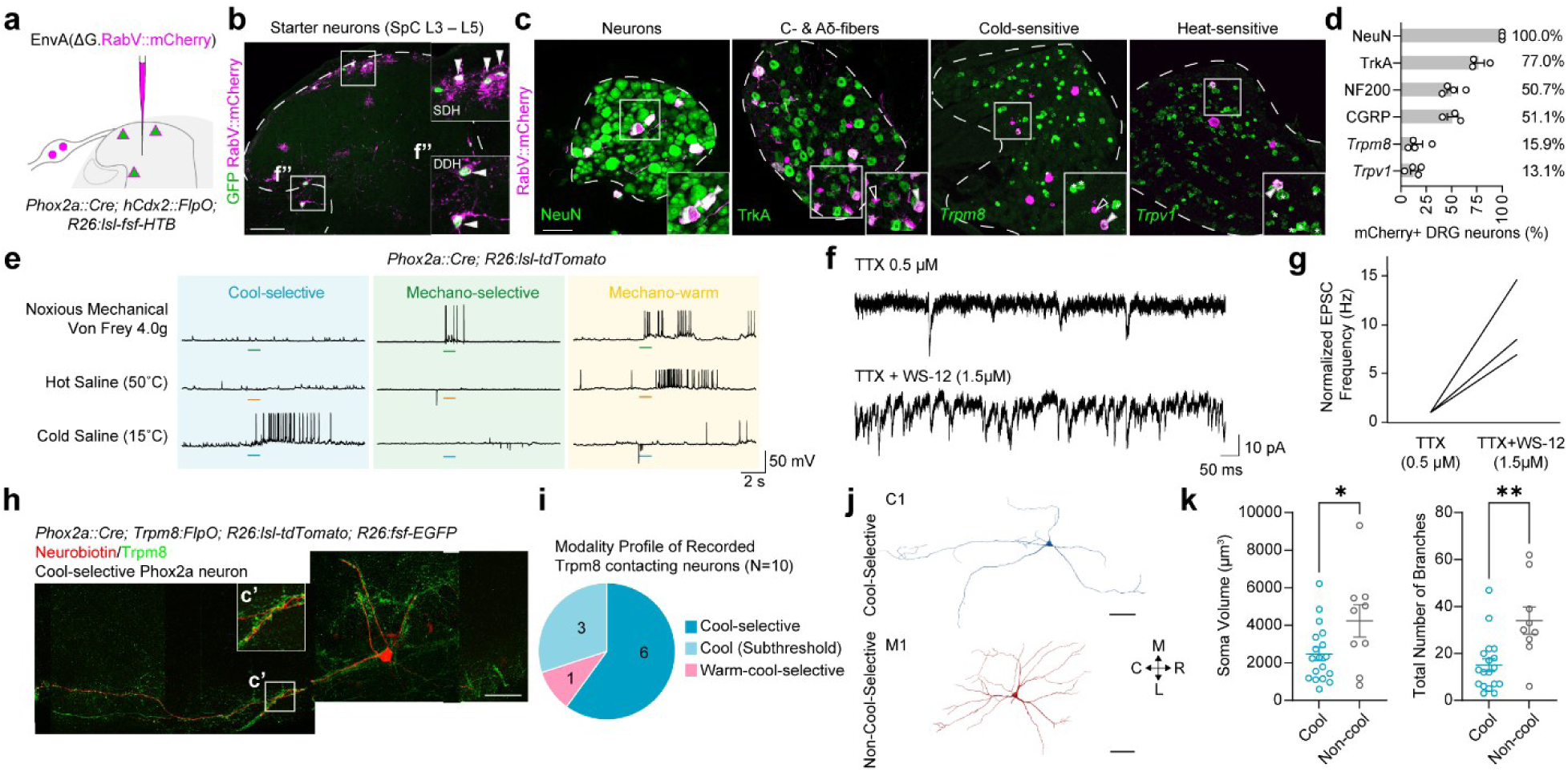
Evidence of thermoreceptor inputs to Phox2a^lamI^ ALT neurons. **a,** Rabies tracing with EnvA(ΔG.RabV::mCherry injection in the lumbar cord of *Phox2a::Cre; hCdx2::FlpO; R26:lsl-fsf-HTB* mice. Spinal Phox2a ALT neurons express histone-GFP and rabies infected cells express mCherry. **b,** Representative transverse spinal cord section containing Phox2a starter cells expressing histone-GFP (green) and mCherry (magenta). **c,** Representative images of DRGs containing rabies infected cells (magenta), expressing sensory neuron marker proteins NeuN or TrkA, or *Trpm8* or *Trpv1* mRNAs (green). White arrow: rabies infected cells expressing the marker, open arrow: rabies infected cells not expressing the marker, *: cells expressing the marker not infected by rabies. **d,** Percentage of rabies infected DRG neurons co-expressing a marker over all rabies infected cells (mCherry+ DRG neurons). **e,** Representative traces of a cool-selective cell (blue), mechano-selective cell (green), or a polymodal cell (yellow) with action potentials evoked by cold saline (15°C, blue), or noxious mechanical (4.0g Von Frey filament, green), or hot saline (50°C, orange). Underbars: stimulus onset. **f, g,** Representative traces **(f)** and normalized EPSC frequency **(g)** of cool-selective Phox2a lamina I ALT neurons following the application of 0.5 μM tetrodotoxin (TTX) abolishing spontaneous EPSCs and reinstated in the presence of 1.5 μM WS-12, a Trpm8 agonist, in addition to TTX (n=3). **h,** Tiled image of a neurobiotin-filled cool-selective Phox2a^lamI^ neuron (red) contacted by Trpm8 fibres (green). **i,** Response profiles of recorded Phox2a^lamI^ neurons (n=10), preselected based on their proximity to Trpm8 fibres, with the majority being cool-selective. **j, k,** Representative reconstructed neurobiotin-filled neurons **(j)** post-recording in **(e)** and somatic and dendritic characteristics **(k)** of the cool-selective cells and non-cool-selective cells. Scale bar: 100 μm in **(b, c, h**) and 50 μm in **(j)**. Mean ± SEM. *p <0.05, **p <0.01, ***p < 0.001, non-significant not labelled. Mann-Whitney test in **(k)**.

We next sought electrophysiological evidence of direct synaptic inputs of Trpm8+ thermoreceptors onto Phox2a^lamI^ neurons, as previous study demonstrated one subpopulation of Phox2a ALT neurons are selectively contacted by Trpm8 fibers^25^. To do so, we conducted whole-cell patch clamp recordings from these neurons in an *ex vivo* semi-intact somatosensory preparation, comprising the continuum of spinal cord, DRGs, saphenous nerve, femoral nerve, and hindlimb skin^30^. Among the 28 Phox2a^lamI^ neurons examined, the majority (19 neurons) responded selectively to cooling (cool-selective), with action potentials elicited by a drop of 15°C saline applied to the skin (skin temperature change: 32°C to 20°C), but not to warming (50°C saline, skin temperature change: 32°C to 38°C) or noxious mechanical stimulation (4.0 von Frey filament, Fig. 3e, Extended Data Fig. 4a). In cool-selective Phox2a^lamI^ ALT neurons (Fig. 3a), spontaneous excitatory post-synaptic currents (EPSCs) were abolished in the presence of tetrodotoxin (TTX) and reinstated in the presence of WS-12, a Trpm8 agonist^31^, in addition to TTX. This implied that the Trpm8+ afferents indeed provide monosynaptic inputs to cool-selective Phox2a^lamI^ ALT neurons (Fig. 3b, c). We further asked whether innervation by cold-selective Trpm8+ thermoreceptors underlies the cool selectivity of Phox2a^lamI^ neurons. To test this, we utilized *Phox2a::Cre; Trpm8:FlpO; R26:lsl-tdTomato; R26:fsf-EGFP* mice to visualize Phox2a^lamI^ neurons and Trpm8 fibers, such that Phox2a neurons expressed tdTomato and Trpm8 afferents expressed GFP (Fig. 4h). Using the *ex vivo* semi-intact preparation, we recorded the thermal responses of Phox2a^lamI^ neurons in close contact with dense Trpm8 fibers. Strikingly, the vast majority (90%) of the these neurons were either cool-selective or exhibited subthreshold response limited to cooling stimuli (Fig. 3i, Extended Data Fig. 4b), suggesting that Trpm8 innervation is likely indictive of cool selectivity in Phox2a^lamI^ ALT neurons.

**Fig. 4.**
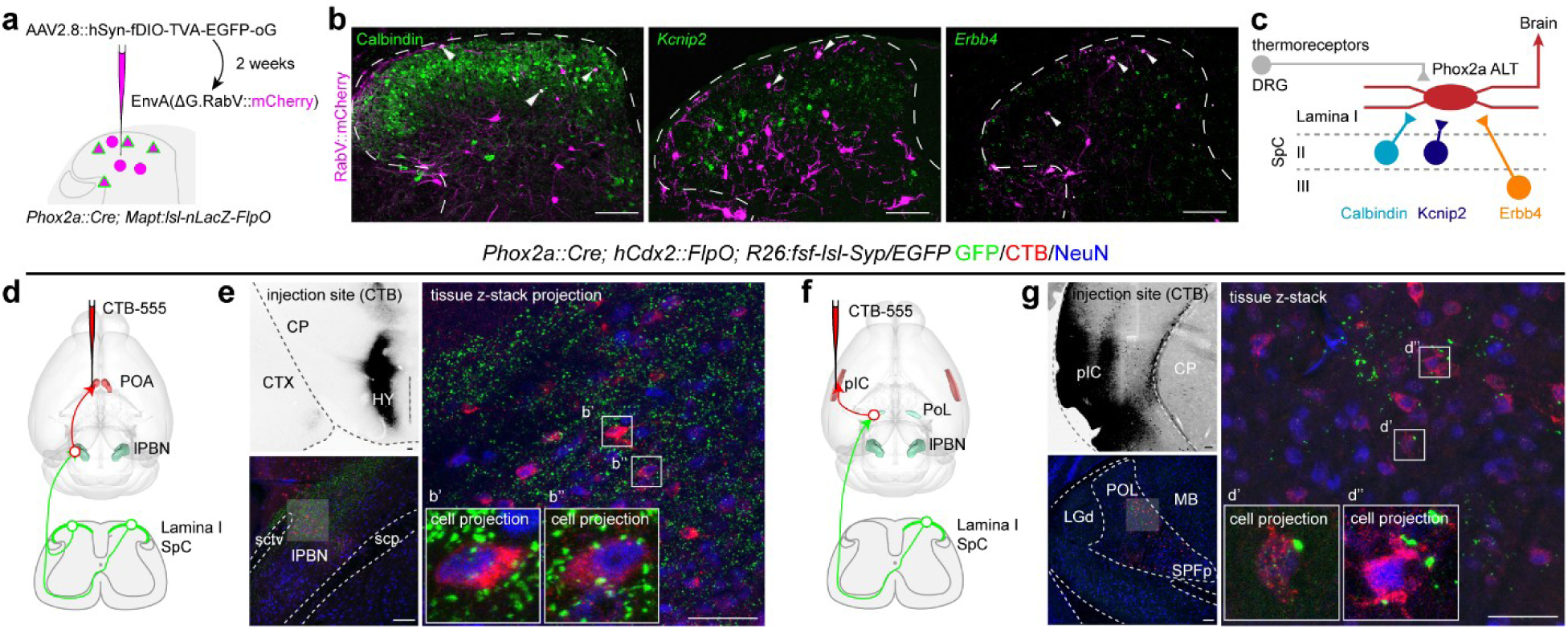
Situating Phox2a ALT neurons in thermosensory circuits. **a,** Monosynaptic rabies tracing experiment via infection with a helper virus and subsequent pseudotyped rabies injected into the lumbar spinal cord. **b,** Rabies-infected spinal dorsal horn neurons (magenta) expressing Calbindin protein, *Kcnip2* and *Erbb4* mRNAs (green). **c,** Summary of identified circuit ofPhox2a^LamI^ neurons. **d,** Experimental design of CTB injection into POA to retrogradely label cell bodies of PBN-POA projection neurons (red) and transgenic expression of EGFP in spinal Phox2a ALT termini (green) in *Phox2a::Cre; hCdx2::FlpO; R26:fsf-lsl-Syn/EGFP* mice. **e,** Axon termini of Phox2a ALT neurons in the lPBN (GFP, green) are juxtaposed on lPBN-POA projecting neuron somata (CTB, red). **f,** Experimental design of CTB labelling of PoL-pIC projections as in **(d)**. **g,** Axon termini in the PoT (GFP, green), on somata of thalamocortical projecting neurons (CTB, red). Abbreviations: CTX (cortex), CP (caudoputamen), HY (hypothalamus), lPBN (lateral parabrachial nucleus), scp (superior cerebellar peduncles), sctv (ventral spinocerebellar tract), pIC (posterior insular cortex), CP (caudoputamen)LGd (dorsal lateral geniculate complex), MB (midbrain), PoL (posterior limiting nucleus of the thalamus), SPFp (subparafascicular nucleus, parvicellular part). Scale bar: 100 μm in **(b, e & g left)** and 50 μm in **(e & g right)**.

In addition, anatomical reconstructions of the 28 recorded Phox2a^lamI^ neurons (diI-filled, from recordings in Fig. 3e) revealed that cool-selective and non-cool-selective (mechano-selective and polymodal) neurons exhibited distinct somatic and dendritic features (Fig. 3j, Extended Data Fig. 4c). The cool-selective neurons had smaller soma volume, and less complex dendritic structure featured with fewer branches and lower density in Sholl analysis, and correspondingly shorter total rostro-caudal and medial-lateral dendritic lengths (Fig. 3k, Extended Data Fig. 4d-f). We also performed principal component-based unsupervised hierarchical clustering (HCPC) of the morphological features of these neurons, which revealed two natural clusters that largely corresponded to non-cool-selective and cool-selective physiological profiles (Extended Data Fig. 4g). Together, these results suggest that Phox2a^lamI^ ALT neurons comprise two main subgroups that are cool-selective and non-cool-selective and correlated with Trpm8 afferent contacts and specific morphological characteristics.

### Situating Phox2a ALT neurons in thermosensory circuits

Beyond peripheral thermoreceptors, thermal information is processed by spinal cord interneurons and brain circuits. First, to assess whether local interneurons provide indirect inputs and modulation onto Phox2a ALT neurons, we performed a new set of monosynaptic rabies tracing experiment as our previous approach yielded insufficient labeling of local interneurons. We utilized an optimized helper virus to express TVA in Phox2a neurons and followed by pseudotyped rabies virus infection (Fig. 4a). Among the rabies-infected cells, we identified Phox2a ALT starter neurons as well as interneurons that are directly presynaptic to them (Extended Data Fig. 5a). The majority of superficial lamina I-III interneurons presynaptic to Phox2a neurons were putatively excitatory, as they did not express inhibitory neuronal marker, paired box 2 (PAX2) transcription factor (Extended Data Fig. 5a, b). This aligns with the high sensitivity of Phox2a^lamI^ neurons to cooling or warming, observed in our calcium imaging result (Fig 2f). Nevertheless, we did observe *Kcnip2*+ (inhibitory) and Calbindin+ (excitatory) interneurons, the two subpopulations previously linked with cooling sensation^32,33^, as well as *Erbb4*+ (excitatory) interneurons associated with heat sensation (Fig 4b)^34^. Thus, in addition to receiving direct inputs from peripheral thermoceptors, Phox2a ALT neurons may also indirectly receive cool and warm temperature-related information from local interneurons (Fig 4c).

We next asked whether supraspinal thermosensory circuits receive direct inputs from Phox2a ALT neurons. Two principal central thermosensory pathways have been proposed: one involves parabrachial neurons projecting to the hypothalamus, primarily the preoptic area (POA), and is implicated in thermoregulatory homeostatic responses. The other includes caudal thalamic neurons, particularly those in the posterior limiting nucleus (PoL), which project to the posterior insular cortex (pIC) for thermal discrimation^10,24^. To determine the relationship between these two pathways and Phox2a ALT neurons, we used *Phox2a::Cre; hCdx2::FlpO; R26:lsl-fsf-SynaptophsinGFP* to visualize axonal terminals of Phox2a ALT neurons (GFP+), and labelled POA-projecting lPBN neurons or pIC-projecting PoL neurons by injecting the retrograde tracer CTB-555 in the POA or pIC, respectively (Fig. 4d, f, Extended Data Fig. 5c). We observed synaptic terminals of spinal Phox2a ALT on the soma of POA-projecting PBN neurons, as well as on pIC-projecting PoL neurons, suggesting directly connectivity between Phox2a ALT neurons and both thermosensory pathways (Fig. 4e, g, Extended Data Fig 5d). Together, our circuit tracing experiments position Phox2a ALT neurons to receive theormosensory information both directly from peripheral thermoreceptors and indirectly from spinal networks, and to channel it to thermoregulatory and discriminative neural pathways.

### Thermoception requires Phox2a ALT neurons

To determine whether Phox2a ALT neurons are functionally required for thermosensation, we selectively expressed diphtheria toxin receptor (DTR) or designer receptors exclusively activated by designer drugs (DREADDs) in spinal Phox2a neurons via the transgenic intersection of *Phox2a::Cre* and *Lbx1:FlpO*. This intersection provided selective access to Phox2a-lineage spinal neurons^35^, capturing 45.6% of lamina I neurons versus 18% of those in the deep dorsal horn (Extended Data Fig. 6a). Moreover, Phox2a neurons in the sympathetic or noradrenergic systems were not captured by this intersection^11^ (Extended Data Fig. 6b), thereby minimizing off-target effects. First, we ablated adult spinal Phox2a neurons by injecting diphtheria toxin (DTX) into *Phox2a::Cre; Lbx1:FlpO; Tau:lsl-fsf-DTR; R26:fsf-lsl-tdTomato* mice (ablated), or into their control (Ctrl) littermates lacking either *Cre*, *FlpO*, or *DTR* transgenes. Two weeks following DTX injection, compared to the control cohort, the number of tdTomato+ neurons was reduced to 10% in the ablated cohort, and their axon terminals in various brain targets were also ablated (Fig. 5a, b, Extended Data Fig. 6c. Moreover, by bilateral retrograde labelling of all ALT neurons from the lPBN, we confirmed that the ablation reduced by approximately half the number of lamina I ALT neurons without significant effects on other spino-lPBN subpopulations (Extended Data Fig. 6d)^16^. Previous studies demonstrated that manipulating thermoreceptor DRG neurons or cool-selective spinal interneurons disrupted temperature discrimination in operant and/or preference task^24,36–38^. Therefore, we asked whether a loss of spinal Phox2a ALT neurons would cause a similar phenotype. In two-plate temperature preference test (TPTP), mice chose between two plates held at either a neutral temperature (30°C) or a test temperature (20°C or 38°C). Compared to the control, ablated mice showed a decreased preference for the 38°C plate (Fig. 5c). Similarly, when mice were placed on a thermal gradient, ranging from 7°C to 52°C, ablated mice showed a less sharp temperature preference peak, and a significant decrease of preference in the 37°C zone compared to control mice (Fig. 5d).

**Fig. 5.**
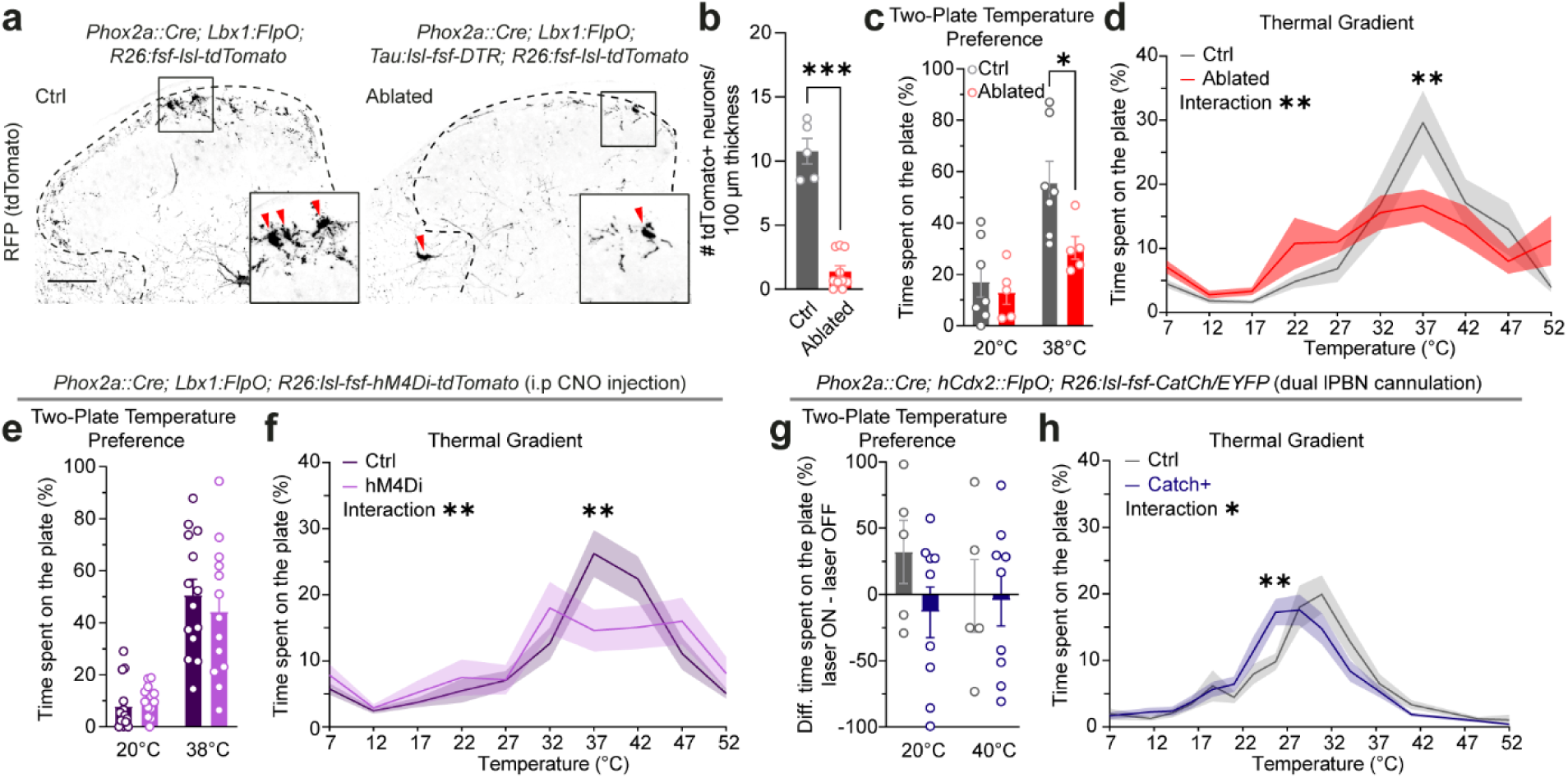
Phox2a ALT neuron function in thermal preference. **a,** Ablation of spinal Phox2a neurons: transverse sections of the lumbar spinal cord from control (Ctrl) or ablated mice showing tdTomato-positive *Phox2a::Cre; Lbx1:FlpO* neurons. **b,** Quantification of tdTomato+ cells in the lumbar spinal cord co-expressing NeuN in control and ablated mice. Cell numbers are averaged per 100 μm thickness (4 sections of 25μm thickness). **c, d,** Behavioural assessments in the control and ablated mice in two-plate temperature preference assay, between 30°C vs. 20°C, or 30°C vs. 38°C **(c)**, and a temperature gradient ranging from 7°C to 52°C and divided into zones of 5°C **(d)**. **e-h,** Behavioural assessments of control and mice with Phox2a ALT neurons being chemogenetically inactivated (hM4Di; **e, f**) or optogenetically activated **(g, h).** Two-plate temperature preference test **(e, g)** was performed as in **(c)** except for difference of percentage of time on the plate was calculated between laser on and laser off in **(g)**. Thermal gradient test **(f, h)** as in **(d).** Scale bar: 100 μm in **(a)**. Mean ± SEM. *p <0.05, **p <0.01, ***p < 0.001, non-significant not labelled. Ordinary Two-way Anova with Bonferroni’s multiple comparisons test in **(c-h)**.

To investigate the effects of acute manipulation of Phox2a ALT neurons, we selectively expressed hM4Di (inhibitory) or hM3Dq (activating) DREADDs in Phox2a ALT neurons (Extended Data Fig. 7a). Thirty minutes following Clozapine N-oxide (CNO) delivery, similar to DTX-ablated mice, hM4Di-expressing mice also showed disrupted preference in the thermal gradient with reduced preference for the 37°C zone (Fig. 5f). In contrast, hM3Dq-expressing mice showed increased preference of the 38°C plate over 30°C in TPTP (Extended Data Fig. 7d).

Therefore, both acute and chronic loss-of-function in Phox2a ALT neurons impaired the thermal preference in mice. Since essentially all Phox2a ALT neurons innervate the lPBN^11,16^, we next examined whether optogenetic activation of Phox2a ALT terminals in bilateral lPBN in *Phox2a::Cre; hCdx2::FlpO; R26:lsl-fsf-CatCh/EYFP* mice would give rise to similar behavioral phenotypes^39^. Like the ablated and chemogenetically-inactivated group, CatCh-expressing mice also showed disrupted thermal preference, shifted towards cooler temperatures with an increased preference in the 25°C zone (Fig. 5h). This suggests that activating warm and cool Phox2a ALT inputs to the lPBN without fine spatial or temporal control may disrupt normal thermal signaling, resembling the effect of input loss. On the other hand, we failed to observe changes in two-plate temperature preference in hM4Di+ and CatCh+ group, or thermogradient preference change in hM3Dq+ group, possibly due to insufficient neuronal activity changes or a plateau effect of preference (Fig. 5e, g, Extended Data Fig. 7e).

In addition to the above measurements that assess the perception of innocuous temperature, we also examined nocifensive responses in the above-described mice models. Ablated mice did not differ in their responses to hot or cold plate (Extended Data Fig. 6f), particularly in the latency to flutter or flinch that are largely rubrospinal-driven, and did not differ in mechanical nociception or locomotion (Extended Data Fig. 6g, h)^21^. In contrast, ablated mice unexpectedly showed increased licking response evoked by both cold stimuli through the application of acetone to the plantar surface of a hindpaw, and intraplantar injection of capsaicin, a TRPV1 channel agonist (Extended Data Fig. 6e). Comparably, we also observed increased capsaicin-evoked licking in the hM4Di group, but not following acetone application (Extended Data Fig. 7b, c), and neither in the hM3Dq+ group (Extended Data Fig. 7f, g). We reason that although Phox2a^lamI^ neurons do not directly encode noxious temperature, they likely coordinate with nociceptive, non-Phox2a ALT neurons to drive nocifensive responses towards noxious thermal stimuli and may modulate analgesia through spinal or descending circuits. Together, our results showed that Phox2a ALT neurons are essential for normal thermal perception, but their contribution to nociception, particularly mechanical, is minimal.

We next considered whether Phox2a ALT neurons also contribute to thermoregulation-impacting metabolic functions, as previously observed in global Trpm8 knockout mice^40^. The metabolic parameters were largely unchanged in the ablated mice (Extended Data Fig. 8h-k), nor did the core body temperature change in ablation, chemogenetic-inactivation, or optogenetic activation (Extended Data Fig. 9e-g). This largely aligns with the observations in studies specifically ablating the peripheral thermoreceptors^41^, indicating that cutaneous thermosensation is dispensable for the gross metabolic function and thermoregulation. However, we did observe an overall increase of loss of water vapor (VH_2_O) along with increased water consumption in the male ablated group, similar to metabolic events during heat dissipation^42^ (Extended Data Fig. 8a-d). Additionally, ablated male mice also displayed higher food consumption and dark-cycle activity (Extended Data Fig. 8e-g). Unexpectedly, although no difference was found in energy expenditure comparing to the control group, male ablated group showed decreased body weight, and lower fat composition (Extended Data Fig. 9a-d). Some of the observed effects could involve the lPBN-POA circuit given its proposed function in thermoregulation, however, other brain targets of Phox2a ALT neurons involved in autonomic functions such as the nucleus tractus solitarius (NTS), parvicellular reticular nucleus (PARN), and intermediate reticular nucleus (IRN) may also participate in thermoregulation^1,43^. Overall, the presence of these metabolic phenotypes in the absence of core temperature shifts and major metabolic parameter changes suggests that Phox2a ALT neurons contribute to nuanced autonomic adjustments that are distinct from the primary homeostatic thermoregulatory response.

## Discussion

The ALT has been extensively studied for its role in nociception, including noxious mechanical and noxious thermal sensation. Although innocuous thermosensitive ALT neurons, particularly cold-selective ones, have also been identified through electrophysiological recordings, their molecular identity and temperature encoding strategies, particularly in innocuous thermal sensing, remain largely unknown^2,7,19^. We uncovered a neuronal ensemble for innocuous thermal perception, genetically defined by the developmental expression of the transcription factor *Phox2a* and comprising around half of lamina I ALT neurons. Our data show that these neurons constitute the central hub of thermosensation within the central nervous system, receiving directly thermal information from peripheral thermoreceptors and dorsal horn thermosensory interneurons, and relaying it to both discriminative thermosensory and thermoregulatory brain pathways.

The existence of central integration and relay of the innocuous thermal information has been implied by recent evidence of large representation of warm and cool neurons in the thalamus and cortex^10,24,44^. Our experiments argue that Phox2a^lamI^ ALT neurons comprise two largely non-overlapping ensembles with preferential tuning to a wide range of warming or cooling temperatures. Given a previous study characterizing all lamina I ALT neurons demonstrated that about half of them responded to cold or warm but not to noxious mechanical stimuli, we infer that the thermal-selective population is likely to be the Phox2a neurons that constitute ∼59% of all lamina I ALT neurons, while the non-Phox2a neurons constitute the noxious mechanical selective population^5,11^. Although *in vitro* and *in vivo* functional studies of thermosensory ALT neurons in rodents and cats described few if any warm-tuned ALT neurons, our calcium imaging experiments identified a surprisingly large cohort of them. Several factors may have contributed to their historic underrepresentation, including the low temporal resolution and sensitivity of Fos induction experiments, absence of descending modulation in *in vitro* electrophysiological recordings^45^, biased sampling of cool vs. warm ALT cells based on the location of antidromic stimulation or fluorescent intensity of tracers used due to soma size differences^10^, and the biased distribution of thermoreceptors in hairy versus glabrous skin in different test conditions^28^.

Our study provides new insights into the mechanism of thermosensation. Notably, cooling-responsive Phox2a^lamI^ neurons showed similar response properties and temporal dynamics as the Trpm8 sensory neurons: both are slow-adapting and encode innocuous cool temperature in a graded manner. Moreover, many cooling-responsive Phox2a ALT neurons and Trpm8 neurons also showed decreased Ca^2+^ responses during warming, known as the “Cool-ON/Warm-OFF” response type mediated by Trpm8 conductance^46^. Therefore, given their monosynaptic connections, the activity of Phox2a cooling-responsive neurons is likely directly dictated by Trpm8-expressing thermoreceptor neurons. Intriguingly, the proportion of innocuous thermal–responsive Phox2a ALT neurons is much higher than that of thermoceptive DRG neurons, particularly the warm ones, suggesting a substantial amplification at the spinal level^4,23,24^. One hypothesis is that besides receiving direct inputs from TRPV1 hot-thermoreceptors, the activation of warm Phox2a ALT neurons may be subject to feedforward control or disinhibition by local circuits^32–34^. Thus, Phox2a ALT neurons comprising the first central nervous system thermosensory node which can integrate and amplify thermal information, instead of merely relaying it.

Prior studies have shown that disruption of periphery thermoreceptors, thalamic nuclei, or the pIC impairs thermal perception^24,36,38^ and our experiments demonstrate a similar requirement for Phox2a ALT neurons. On the other hand, although manipulating brain thermoregulatory circuits that receive ALT-relayed information resulted in abnormal thermoregulation^47,48^, neither peripheral thermoreceptor deletion, nor our Phox2a ALT ablation resulted in notable thermoregulatory or metabolic defects^41,49^. Thus disrupting cutaneous thermosensation or its relay may be compensated by redundant thermoregulatory pathways involving airways, autonomic nervous system, and/or brown adipose tissue^50^. More generally, our experiments support a model in which discriminatory thermal perception and thermoregulation are processed in parallel and independently^10^.

In sum, our work provides a foundation for understanding of neural mechanisms of thermal perception, and thermal pain hypersensitivity in pathological conditions. The evolutionary conservation of thermal encoding strategies and Phox2a expression, suggests the transmission of innocuous thermosensory information by cool and warm selective ALT neurons may be part of a global molecular logic of thermosensation^3,8,10,11,45^.

## Methods

### Mouse line generation

All procedures were approved by local committees, including the IRCM and Université Laval Animal Care Committee, using Canadian Council for Animal Care (CCAC) regulations, and the Ethical Review Process Applications Panel of the University of Glasgow, according to the UK Animals (Scientific Procedures) Act 1986 and ARRIVE guidelines. Adult male and female mice (6-19 weeks of age) were used, and sex ratio for all experiments was kept close to 1:1. Mice were housed with a 12-hour: 12-hour light: dark cycle (light: 6h00 to 18h00) with food and water *ad libitum*. Mice were maintained on a mixed 129/Sv and C57BL/6J, or pure C57BL/6J background as indicated in the key resource table. In the ablation, chemogenetic activation, chemogenetic inactivation, and optogenetic behavioural experiments, mouse lines containing *Phox2aCre/+* and *Lbx1FlpO/+* or *hCdx2FlpO/+* was first generated^35,51^, and bred with parents containing *Tau:DS-DTR-/-, Ai65-/-* (ablation), or *Rosa:DS hM3Dq-mCherry-/-* (chemogenetic activation), or Rosa:DS-hM4Di-tdTomato-/- (chemogenetic inactivation), or *Rosa:DS-CatCh-/-* (optogenetic) to generate experimental mice (Phox2a^abl^, Phox2a^hM3Dq^, Phox2a^hm4Di^, or Phox2a^CatCh^) that carry all transgenes, and littermate control mice that lack either *Phox2aCre* or *Lbx1FlpO* transgenes. For detailed information of mouse strains used, see Supplementary Table 2.

### Immunohistochemistry

Tissue dissection was as previously described^11,52^. For adult tissue collection, mice were first anesthetized and transcardially perfused with 1xPBS solution followed by 4% PFA. Brains,spinal cords and DRGs were collected and post-fixed overnight in 4% PFA on a shaker at 4°C, followed by overnight wash in 1x PBS, and transfer to 30% sucrose until sunk. For embryonic tissue collection, vaginal plugs were checked daily at 6h00, and the day of plug detection was noted as embryonic day 0.5 (E0.5). On the day of E16.5, mothers were anesthetised with isoflurane and received a cervical dislocation. Embryos were dissected in 1xPBS and followed by the same fixation procedure as described above. Tissue was embedded in O.C.T. and stored at -80°C until cryosectioning mounting on Superfrost plus slides which were dried and stored at -80°C until processed for immunofluorescent protein or mRNA detection. For the former, slides were rehydrated and washed in 1xPBS, and blocked with blocking solution (5% Normal Donkey Serum (NDS) in 1xPBST (0.1% Triton X-100 in 1xPBS)) for 30 minutes at room temperature (RT). Primary antibodies were added in 1% NDS blocking solution and coverslipped to incubate overnight at 4°C. Slides were then washed with 1xPBS, and the secondary antibodies in 1xPBST solution was applied for 2 hours at RT. Finally, the slides were washed in 1xPBS, mounted with Mowiol, and imaged upon dried. For all detailed reagents used, see Supplementary Table 2.

### *In situ* mRNA localisation

Cryosectioned slides were processed using RNAscope Multiplex Fluorescent v2 kits as per manufacturer’s protocol (https://acdbio.com/manual-assays-rnascope). For some Figure 1 and Figure S1 experiments indicated in figure legends, immunohistochemical protein detection was performed as above following mRNA detection and prior to the mounting.

### Stereotaxic surgery

Mice received 1 mg/kg of body weight of buprenorphine intraperitoneal (i.p.) the day of the surgery. During the surgery, mice were anesthetised with 4% isoflurane (induction phase) followed by 2% isoflurane (maintenance phase) in a mixture of 70% N_2_ and 30% O_2_, and placed in a stereotaxic apparatus. Following shaving, the skin was disinfected with iodine and ethanol. Intracranial and intraspinal injections were performed generally as previously described^11,53^. For all intracranial injections, the following coordinates (in mm) and volumes for single injection of CTB were used: 5.0 AP, ±1.5 ML, 2.75 DV, 250 nL for PBN, 0.5 AP, -4.25 ML, 3.0 DV, 300 nL for pIC, 0.1 AP, -0.3 ML, 5.8 DV, 300 nL for POA, similar to those previously described^11^. For intraspinal injections, mice were anesthetised as above, and the intervertebral space between T13 and L1 of vertebral bones (L3-L4 lumbar spinal cord) was exposed. Three injections of 333 nL AAVs or rabies viruses were made along the AP axis (spaced 400 µm apart along the AP axis, 0.48 ML, 0.26 DV)^54^. All injections were performed at a rate of 100 nL/minute, and the glass capillary was held for another 3 minutes after the injection to prevent leakage. Mice recovered in a heated chamber until ambulatory before returning to their home cages.

### Patch-clamp recording in semi-intact preparation

Semi-intact preparation was performed largely as previously described^19,30^. Briefly, adult mice were perfused with oxygenated (95% O_2_/5% CO_2_) sucrose-based artificial cerebrospinal fluid (ACSF) (in mM; 3.0 KCl, 1.2 NaH_2_PO_4_, 0.5 CaCl_2_, 7.0 MgCl_2_, 26.0 NaHCO_3_, 15.0 glucose, 251.6 sucrose, 1.0 Na ascorbate, and 1.0 Na pyruvate) at room temperature. Immediately after perfusion, the spinal cord (∼C2-S6) and the right hindlimb were excised and submerged in sucrose-based ACSF circulated at 5 mL/min in a Sylgard-lined dish. Next, the saphenous nerve and the femoral cutaneous nerve and the innervated skin were dissected, and the L2 and L3 dorsal root ganglion were kept on the spinal cord. Dural and pial membranes were carefully removed and spinal cord was pinned onto the Sylgard block with the right dorsal horn facing upward. The chamber was then transferred to the rig and rinsed with normal ACSF solution (in mM; 125.8 NaCl, 3.0 KCl, 1.2 NaH_2_PO_4_, 2.4 CaCl_2_, 1.3 MgCl_2_, 26.0 NaHCO_3_, and 15.0 glucose) saturated with 95% O2/5% CO2 at 25°C for at least 15 minutes. Patch-clamp recordings were performed up to 4 hours after dissection. Neurons were first visualized with a fixed-stage upright microscope and a narrow-beam infrared LED as described previously. Whole-cell patch-clamp recordings were made with a pipette constructed from thin walled single-filamented borosilicate glass with resistances of 6 -12 MΩ. Electrodes were filled with an intracellular solution (in mM; 130.0 K gluconate, 10.0 KCl, 2.0 MgCl_2_, 10.0 HEPES, 0.5 EGTA, 2.0 ATP-Na, 0.5 GTP-Na, and 0.2% Neurobiotin, pH adjusted to 7.3 with 1.0 M KOH). Neurobiotin (Vector Labs Peterborough, UK) was added to confirm recording from the target cell. Signals were acquired with an amplifier (Axopatch 200B; Molecular Devices, Sunnyvale, CA). The data were low-pass filtered at 2 kHz and digitized at 10 kHz with an A/D converter (Digidata 1550B; Molecular Devices) and stored using a data acquisition program (Clampex version 11; Molecular Devices). The liquid junction potential was not corrected. Mechanical stimulation was applied using a 4.0 g von Frey filament. Thermal stimulation was applied using hot (50°C) and cold (15°C) saline applied gently to the skin using an eye dropper.

### Morphological analysis

Neurobiotin-labelled cells from electrophysiological recordings were revealed by incubating in avidin-Alexa 555 (1:1000; Jackson ImmunoResearch) in 1xPBS containing 0.3% Triton X-100. They were imaged with a confocal microscope through a 63× oil-immersion lens (numerical aperture 1.4) at 0.5-μm z-spacing with all visible and distinguishable axonal and dendritic arbors. Morphometric data for dendritic trees of all the reconstructed cells were obtained from Neurolucida Explorer. The dendritic parameters were extracted as described previously^55^. To make an objective comparison between the cool-selective and non-cool selective neurons, we performed cluster analysis using Ward’s method as described previously^55,56^. To reduce the dimensionality of the original data while preserving variance, we calculated principal components from the data set. The number of principal components to be retained for cluster analysis was then determined from a screen test.

### *In vivo* calcium imaging

*In vivo* Ca2+ imaging in *Phox2a::Cre; R26:lsl-GCamP6s* mice was performed largely as described previously^23^. Briefly, animals were anesthetised with 100 mg ketamine, 15 mg xylazine, and 2.5 mg acepromazine per kg of body weight in saline, and then received a lumbar laminectomy and were fixed on a spinal stabilizing apparatus. L3-L4 region was imaged with a two-photon microscope using a 940 nm tunable femtosecond laser. RAW image sequences were acquired at 1000 x 500 pixels at a rate of 32Hz and with a pixel size of 0.385 μm. During the imaging, the plantar side of the hind paw of the animal was stimulated by repeated manual brushing with a paintbrush, pinching with serrated forceps, or thermal stimuli delivered by a feedback-controlled Peltier device and a 1 x 1 cm thermal probe (TSA-II-NeuroSensory Analyzer, Medoc). Each stimulus was provided with at least 3-5 minutes intervals and started with innocuous brush and thermal stimuli, then noxious pinch to prevent sensitization. After acquisition, RAW image sequences were converted into TIFF format in ImageJ and registered with a custom-built MATLAB (MathWorks) function through rigid body translation alignment based on the 2D cross-correlation to correct for movement. Background pixel value was defined by the average pixel value of a region of interest (ROI) containing no visible neurons. Neuronal ROIs were drawn manually in the cytoplasm of visible neurons. The average fluorescence intensity of a given ROI was defined as Ft, calculate as the mean pixel intensity within that neuronal ROI minus that of the background ROI. Ca2+ amplitude was defined as ΔF/F0 = (Ft-F0)/F0, where F0 is the baseline fluorescence value, measured as the average of the first two seconds of Ft. Positive responses were detected and measured automatically using a custom tool written in Spike2 (Cambridge Electronic Design [CED]) and confirmed by visual inspection, and all data were exported into Microsoft Excel Table for subsequent statistical analysis^23,57^.

### Behavioural assays

The experimenters were blinded to genotypes during the experiment and video analysis. Prior to all tests, mice were habituated to the behaviour test room for at least 30 minutes in their home cages except for von Frey and acetone test when mice were habituated in the test chambers, and test chambers and apparatus were always cleaned and deodorized with Peroxygard between animals. Behaviour tests except vF were filmed for post hoc analysis. All tests were performed on separate days to prevent sensitization or adaptation.

The **thermogradient test** was performed using Thermal Gradient Test 2.0 (Bioseb)^58^. In brief, a 120 cm metal corridor was set at a continuous and stable temperature gradient from 7°C to 52°C, and its orientation was randomized. Ten virtual temperature zones with a 5°C-step difference at the centre were defined for two lanes for two mice for each test. Mice were introduced in the middle of the corridor and allowed to explore freely for 30 minutes, and their position was recorded by an overhead camera for video tracking. The percentage of cumulative time spent in each temperature zone was reported for the second half of the test.

The **two-plate temperature preference test** was performed with Thermal Place Preference-Two Temperatures Choice Nociception Test (Bioseb)^58^. Mice were allowed to explore the two compartments of the metal plate set at experimental temperatures (38°C or 20°C) and neutral temperatures (30°C) for 10 minutes with overhead camera recording for video tracking. Mice were first tested for their preference with both plates set at 30°C (Day 1), and those exhibited high spatial preference (>75% of one compartment) were excluded from the analysis. Mice were then tested for their preference of the experimental temperature on Day 2 and Day 3, with the sequence of the temperature and its compartment randomized.

For the **von Frey test**, mice were placed in a plexiglass chamber (4 × 2.2 × 2.5 cm each) with metal dividers on a mesh floor. A set of nylon von Frey filaments (0.008, 0.02, 0.04, 0.07, 0.16, 0.4, 0.6, 1.0, 1.4 g) were used to stimulate the plantar surface of the mouse hind paw, using the “up-down” method of Dixon, and withdraw threshold was calculated accordingly, ranging from 0 (most sensitive) to 2 (least sensitive)^59^.. The filaments were calibrated every time prior to the test.

The **foot clip test** was performed as performed as described previously^60^. Briefly, mice were pinched on the hindpaw with a toothless mechanical clip and then placed into a plexiglass cylinder on a plexiglass sheet and video recorded from below for 60 seconds. The total amount of time licking the clipped paw was recorded.

For the **hot plate and cold plate test**, mice were placed in the hot-cold plate apparatus (4” x 8”, IITC PE34) at set temperature of 53°C ± 0.1°C (hot plate) or 1°C ± 0.1°C (cold plate), and video recorded from front-above with a mirror at the back of the chamber to provide clear view of both sides of the mice^11,61^. For the hot plate, mice were removed once licking of both hind paws were observed, or cutoff at 15 seconds. For the cold plate, mice were recorded for 60 seconds. The latency of withdraw of both paws were recorded immediately during the test, and the videos were further analysed to record the latency to lick the hind paws. Scores were arbitrarily assigned to the cold plate test as the following: 0 – no reaction, 1 – tiptoeing or avoidance, 2 – withdrawal or flinching, 3 – licking.

For the **acetone test**^11,62^ mice were placed in a plexiglass chamber (4 × 2.2 × 2.5 cm each) with metal dividers on a mesh floor. A drop of acetone extruded from the blunt end of a 5ml syringe was applied to either hind paw of the mouse. Mice were stimulated 5 times in total, with an inter-trial-interval of 5 minutes, and each trial was recorded for 60 seconds following the stimulation. The videos were analysed with BORIS and the total licking time was reported as a sum of 5 trials^63^.

For the **capsaicin test**^11,61^, 20 μL capsaicin solution (1.5 ug/20 ul in 1xPBS) at RT was injected intraplantarly in the hind paw using a standard 28G insulin syringe (BD). Mice were placed in a plexiglass cylinder with blocked visual to other mice. The video was recorded for 15 minutes and then analysed for the duration of licking behaviour.

For **the open field test**, mice were placed in a square chamber (50 x 50 x 40 cm) and allowed to freely explore for 30 minutes^64^. An overhead tracking camera recorded their movement in the chamber and the cumulative distance they travelled was reported.

### Optogenetic Stimulation

Animals were implanted with optical fibers (NA 0.22, 200 μm core, Doric) over the lateral parabrachial nucleus (AP: -5.20, ML: ±1.40, DV: -2.50, based on The Mouse Brain in Stereotaxic coordinates, 2nd edition, ISBN 0-12-547636-1, Paxinos & Franklin^65^). The laser intensity was 10mW on each side of 488 nm light (20 Hz, 5 ms-duration pulses). Mice were habituated to the cables prior to behavioural tests.

### Quantification and statistical analysis

Confocal images of cells are processed with ImageJ software and are quantified with its cell counter plugin. Video recordings of mice behaviours are quantified using Behavioral Observation Research Interactive Software (BORIS) and annotated frame based. Data was recorded and sorted in Microsoft Excel and imported in GraphPad Prism software for statistical analysis and display. Significance is represented as ns: nonsignificant, *p < 0.05, **p < 0.01 or ***p < 0.001 for all figures.

## Data availability

All other data are available in the main paper and supplementary materials. All materials are available through requests to the corresponding author.

## Acknowledgements

We thank the Goulding lab for sharing the following mouse lines with us: *hCdx2::FlpO; Rosa26:ds-HTB; Lbx1::FlpO; Tau:ds-DTR*, Dr. Mark Hoon for sharing *Trpm8:Flp* mice, Dr. Marie-Eve Paquet and her team for developing all the AAVs and Rabies viruses used in this project (Viral Vector Core of Canadian Neurophotonics Platform), Dr. Magali Millecamps (ABC Platform, McGill Univeristy) for contributing to the thermogradient, two-plate temperature preference, light brush, and hot/cold plate tests, Meirong Liang for genotyping. This work was supported by project grants from the Canadian Institutes of Health Research (CIHR; PJT-162225, MOP-77556, PJT-153053, PJT-183824, PJT-159839, and PJT-197987) to A.K., Natural Science and Engineering Research Council Discovery grant CG139292 to F. W., and the Medical Research Council (MR/V033638/1) to J.H. and A.J.T. A.K. held the Doggone Foundation Chair of Excellence in Pain and received support from the Foundation IRCM. X.Z. is a recipient of a Fonds de recherche du Québec -Santé doctoral scholarship. We thank Drs. Allan Basbaum, Brian Roome, Bo Duan, Ran Chen, and Sebastian Choi for their generous feedback on the manuscript and the project.

## Author contributions

X. Z. Performed all experiments except for the electrophysiology, calcium imaging and partial behavioural experiments as described in Methods, and analysed the data. Partial behavioural experiments were performed by M.M and S.F., and analyzed by X.Z. F.W. performed the calcium imaging experiments. A.M., M.K., and J.H. performed the electrophysiological experiments, and K.A.B. and A.M. performed subsequent morphological analyses, M. E. P. provided key reagents, A.K., Y.d.K., J. H. and A.J.T. supervised the study and acquired funding. A.K. and X.Z. wrote the manuscript.

## Extended Data

**Extended Data Fig. 1.**
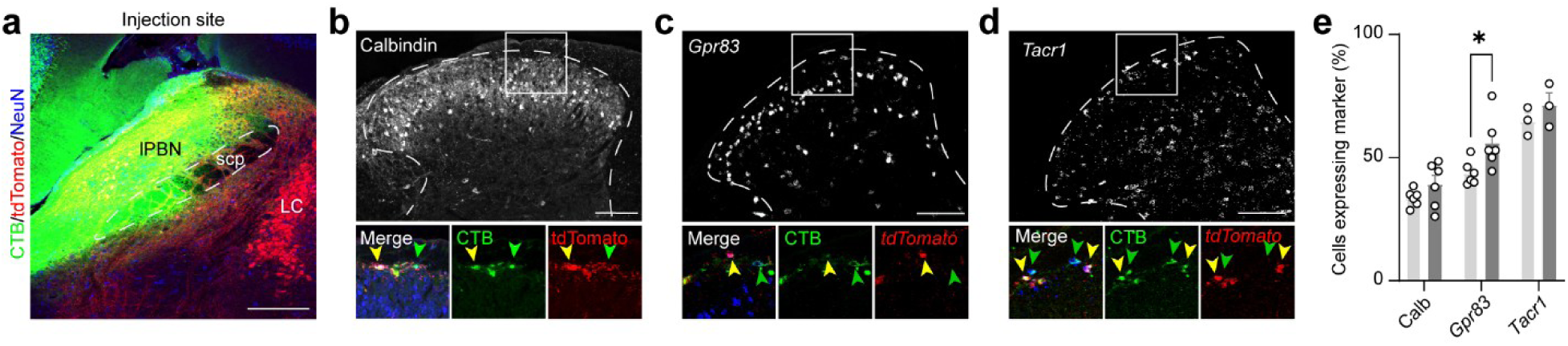
Additional characterisation of lumbar Phox2a ALT neurons. **a,** Representative images of the injection site in the lateral parabrachial nucleus (lPBN). **b-d,** Additional staining of CTB (green), tdTomato (red), and Calbindin antibody (white, **b**) or RNAscope probes targeting tdTomato (red) and Gpr83 **(c)** or Tacr1 mRNA (white, **d**) in Phox2a+ (yellow arrowhead) or Phox2a- (green arrowhead) ALT neurons. **e,** Quantification of the percentage of Phox2a+ (light gray) or Phox2a- (dark gray) ALT neurons co-expressing the marker. Scale bar: 100 μm in **(a-d)**. *p <0.05, **p <0.01, ***p < 0.001, non-significant not reported. Mann-Whitney test in **(e)**.

**Extended Data Fig. 2.**
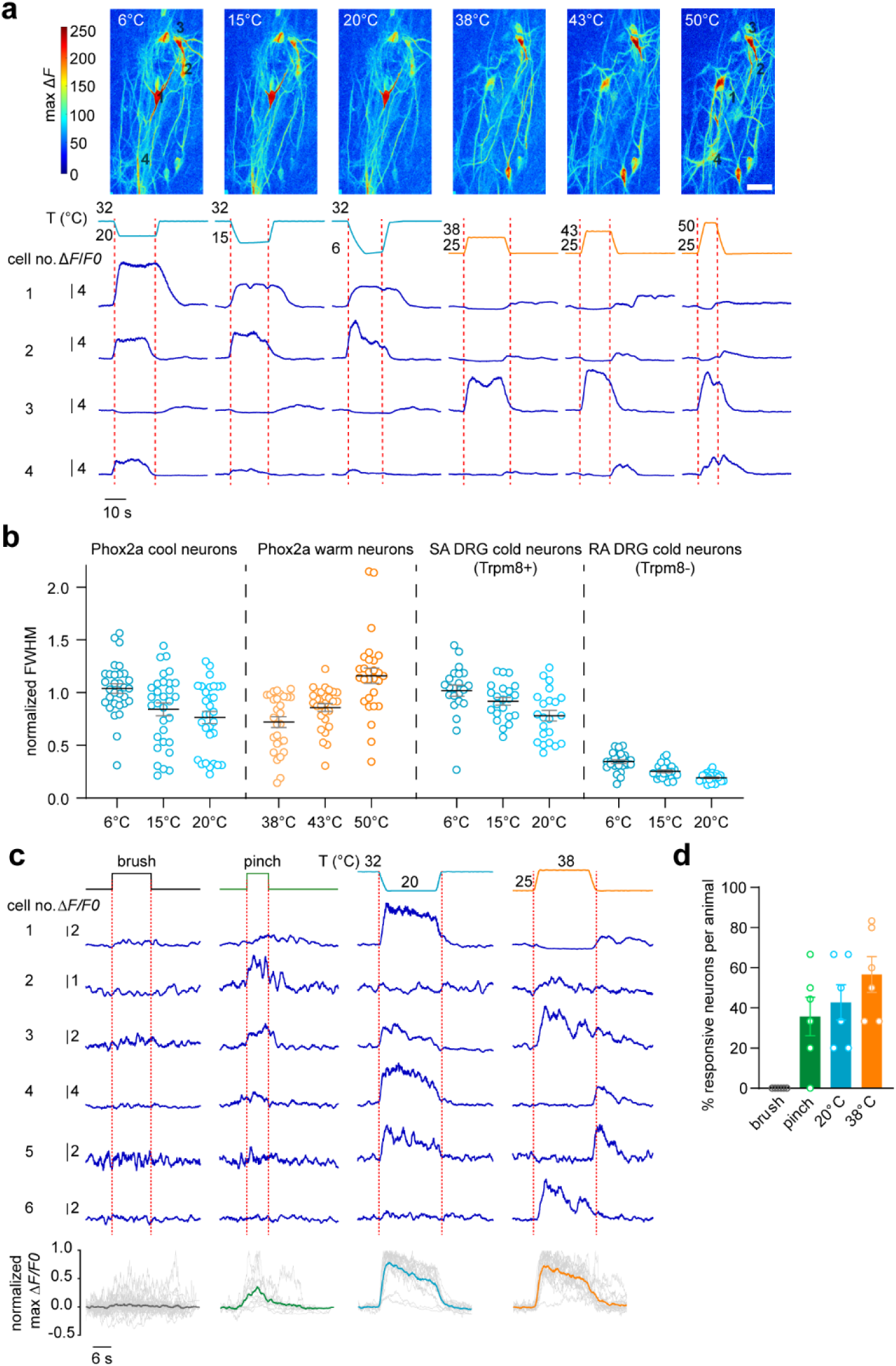
Phox2a lamina I ALT neurons are tuned to temperature changes. **a,** Representative imaging window and calcium curves from four neurons showing typical responses to thermal stimuli as in Figure 1 **(h**), cell number same as the row number. Cell number 1 and 2 are typical cool-selective Phox2a neurons, cell number 3 is a typical warm-selective Phox2a neuron, and cell number 4 is a cold-heat responding Phox2a neuron. **b,** Normalized full-width-half-maximum (FWHM) of calcium responses over thermal stimulation duration in Phox2a cool-selective or warm-selective neurons showing a slow adaptive (SA) characteristic similar to SA cool (Trpm8+) thermoreceptors but distinct from rapid adaptive (RA) cool thermoreceptors. **c, Top:** representative calcium curves from six neurons in response to manual brush, manual pinch, or thermal stimuli as in Fig. 1 **(h)**. Bottom: individual calcium traces and the population average from all neurons tested. **d,** Summary of the percentage of neurons responded to each stimulus within each animal (N=6).

**Extended Data Fig. 3.**
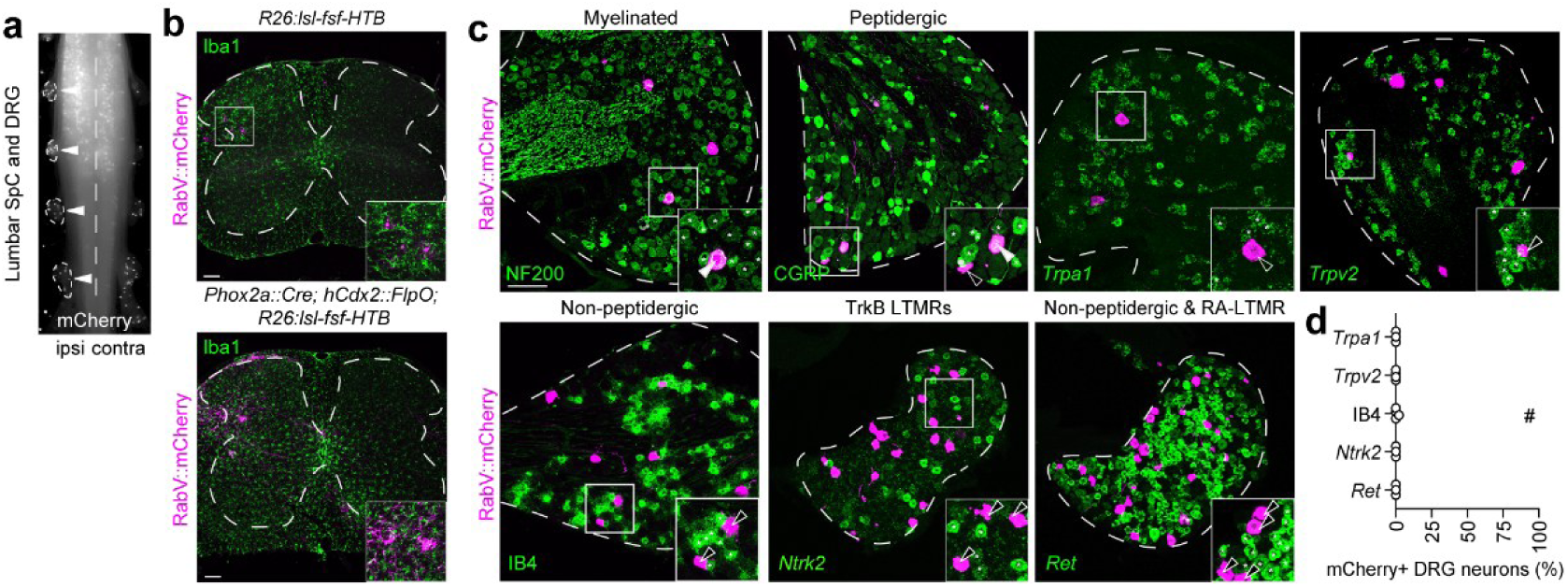
Sensory neuron inputs to Phox2a ALT neurons. **a,** Representative image of whole-mount lumbar spinal cord with DRGs attached containing rabies infected cell expressing mCherry (white). **b,** Representative images showing that control animals with no Cre and FlpO (TVA-) were not infected by rabies in neurons, but rabies infected microglia in both TVA- and TVA+ animals. **c,** Representative images of rabies-infected neurons not co-expressing Ntrk2, Ret, Trpv2 mRNA or IB4 protein. **d,** Quantification of the percentage of rabies infected DRG neurons co-expressing the markers over all rabies infected cells (mCherry+ DRG neurons). #: rabies-resistant non-peptidergic neurons expressing IB4. **e,** Cell body sizes of all rabies-infected DRG neurons. Both small, median, and large diameter neurons were found., but the majority are small to median size. Each curve represents an individual animal, N=5. Scale bar: 100μm in **(b, c)**.

**Figure S4.**
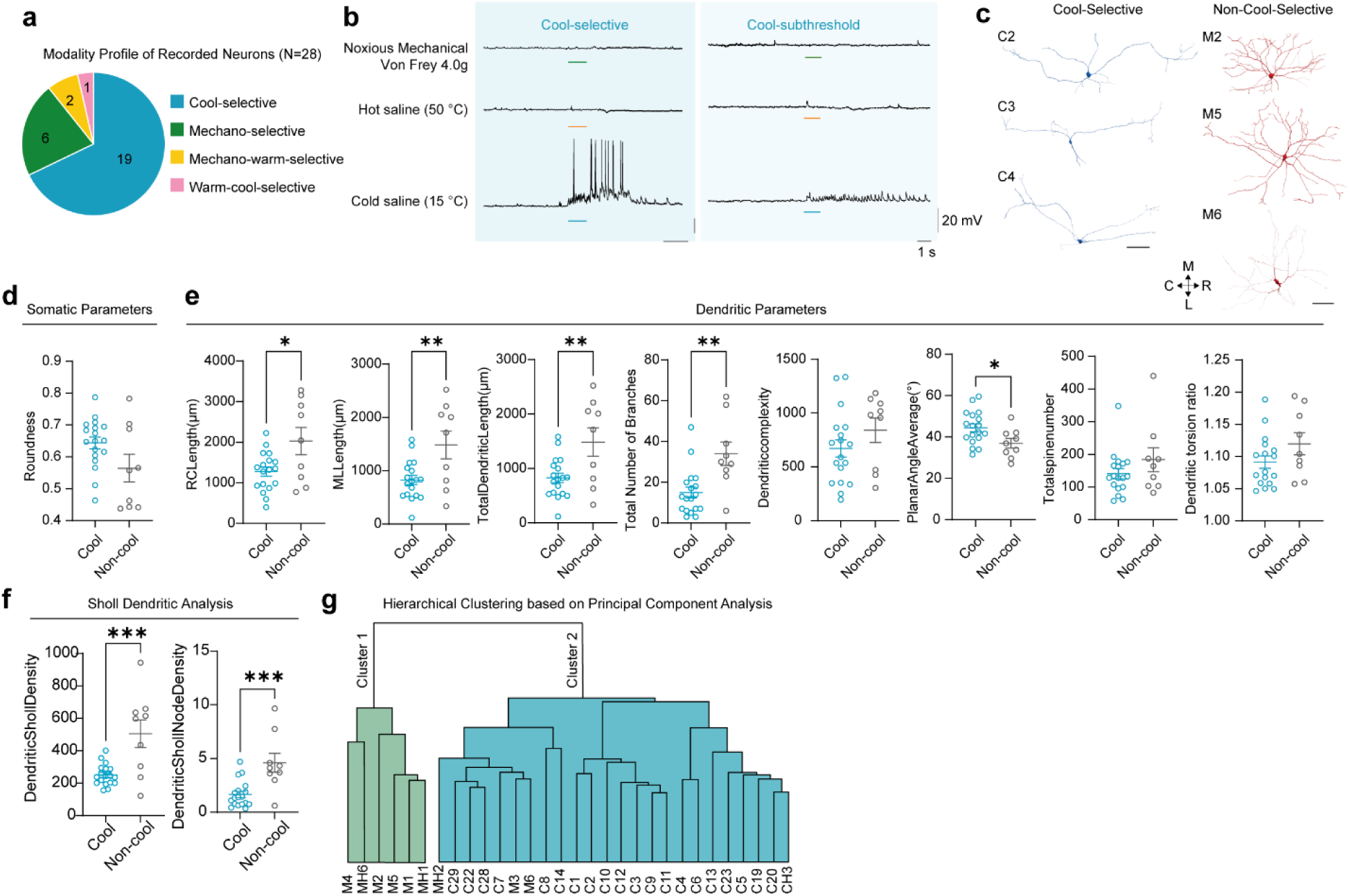
Further physiological and morphological characterization of Phox2a^LamI^ ALT neurons. **a,** Pie chart of the response profile to all modalities of stimuli of all Phox2a^lamI^ neurons recorded (N=28). **b,** Representative traces of cool-selective and cool-subthreshold Phox2a^lamI^ cells with Trpm8 fibre contact, similar as described in **(Fig. 3e)**. **c,** Additional reconstructed cells of cool-selective and non-cool selective Phox2a^lamI^ neurons. **d-f,** Other somatic and dendritic parameters of cool-selective and non-cool selective Phox2a^lamI^ neurons. **g,** Two major clusters identified by hierarchical clustering based on the principal component analysis of the morphological parameters from neurobiotin-filled cells. C: cool-selective, M: mechano-selective, MH: mechano-warm-selective, CH: warm-cool-selective. *p <0.05, **p <0.01, ***p < 0.001, non-significant not reported. Mann-Whitney test in **(d-f)**.

**Extended Data Fig. 5.**
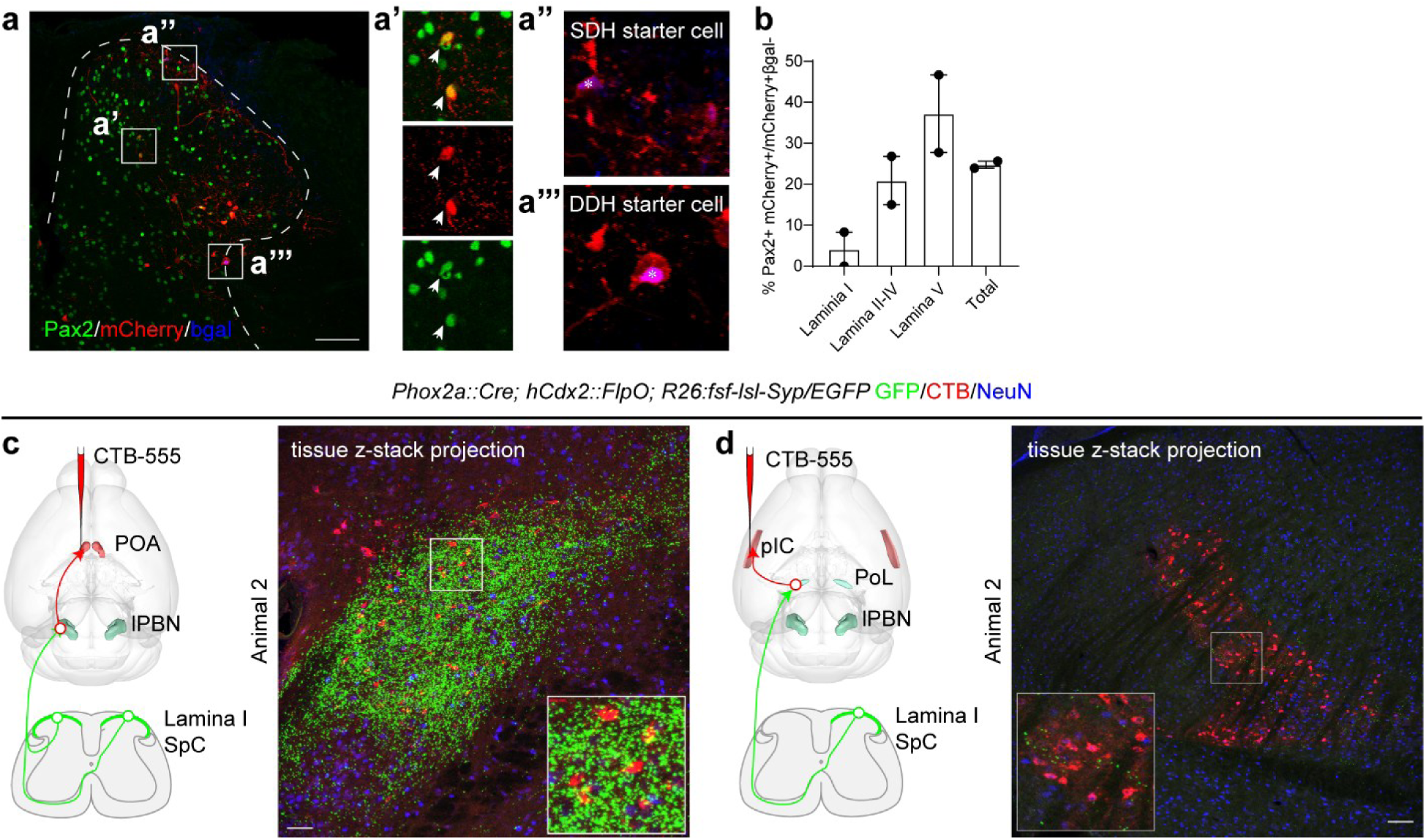
Additional characterization of Phox2a neurons in local and supraspinal circuits. **a,** Representative picture of local spinal dorsal horn infected with rabies (mCherry, red) and labeling of the inhibitory neuron marker Pax2 (green). Starter cell revealed by β-galactosidase expression (blue, **a’’** and **a’’’**). **b,** Percentage of interneurons presynaptic to Phox2a neurons that are inhibitory (Pax2+), shown by laminar distribution. **c,** Additional animal showing Phox2a axon terminals on lPBN-POA projecting neurons, observed in total N=5 animals with similar pattern**. d,** Additional animal showing Phox2a axon terminals on PoL-pIC projecting neurons, observed in total N=3 animals with similar pattern. Scale bar 100 μm in all.

**Extended Data Fig. 6.**
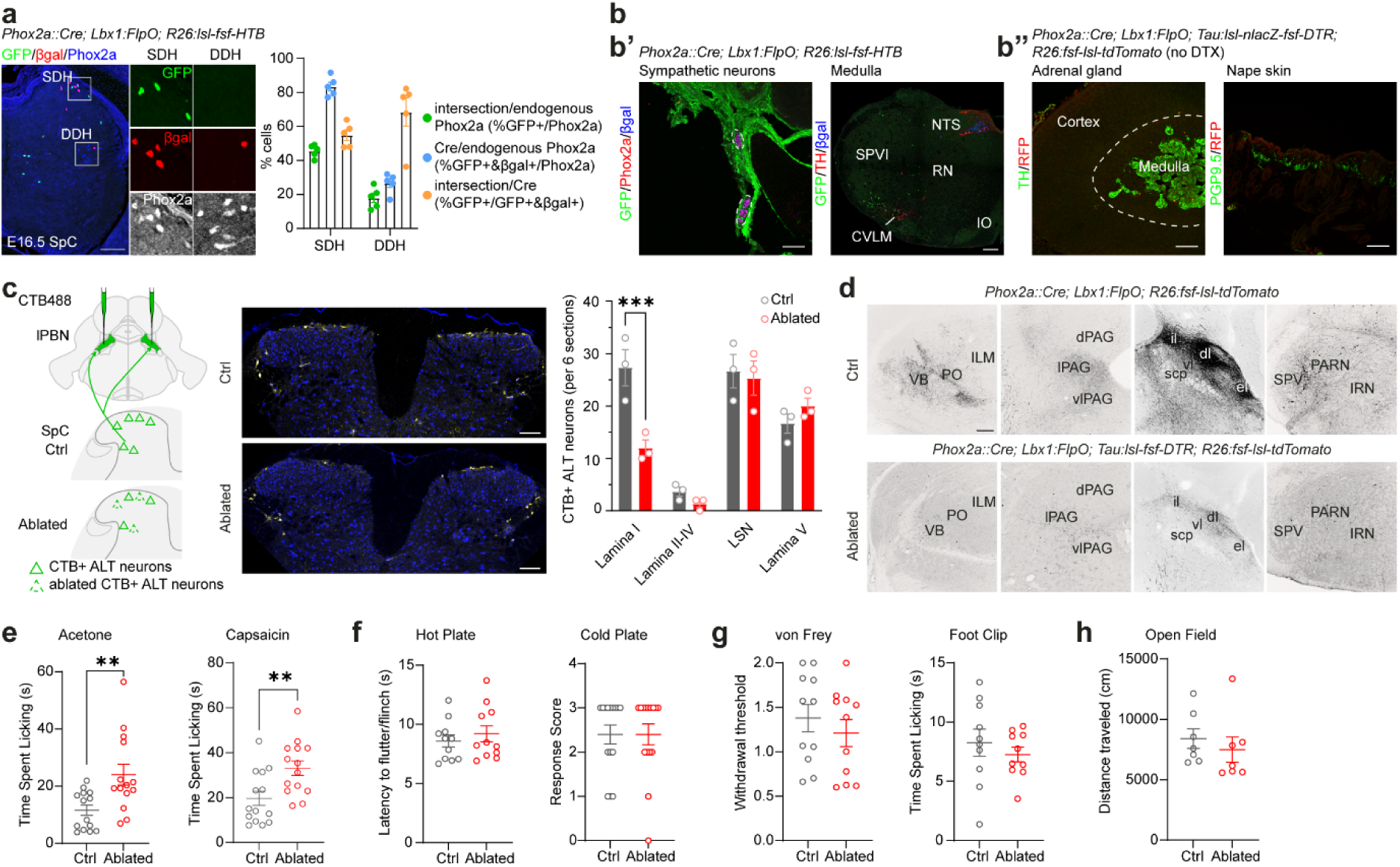
Characterization of the ablation model and nocifensive and locomotor behaviours. **a,** Intersectional efficiency of *Phox2a::Cre* alone or with *Lbx1:FlpO* in the superficial dorsal horn (SDH) and deep dorsal horn (DDH) in comparison to the endogenous Phox2a protein expression. β-galactosidease: Cre expression only; GFP: Cre and FlpO co-expression. **b,** *Phox2a::Cre; Lbx1:FlpO* intersection does not target cells in the sympathetic ganglion, noradrenergic (TH+) medullary areas, adrenal gland, or skin at the nape area. **c,** Representative pictures showing that the ablation mainly decreased the number of lamina I but not deep laminae ALT neurons (CTB, green). **d,** Ablated animals showed loss of innervation in many brain areas that normally receive abundant Phox2a ALT innervations as in control (ctrl). **e-h,** Licking responses to acetone and capsaicin application were increased in Ablated group **(e)**. Reflexive withdrawal in response to hot and cold plate **(f)**, as well as to von Frey filaments stimulation (g), licking responses to mechanical stimuli **(g)**, and general locomotion **(h)** did not differ between Ablated and control animals. Scale bar: 100 μm in **(a, b’, c)**; 200 μm in **(b’’, d)**. Mean ± SEM. * p<0.05, ** p<0.01, *** p<0.001. Mann-Whitney test **(e-g)**.

**Extended Data Fig. 7.**
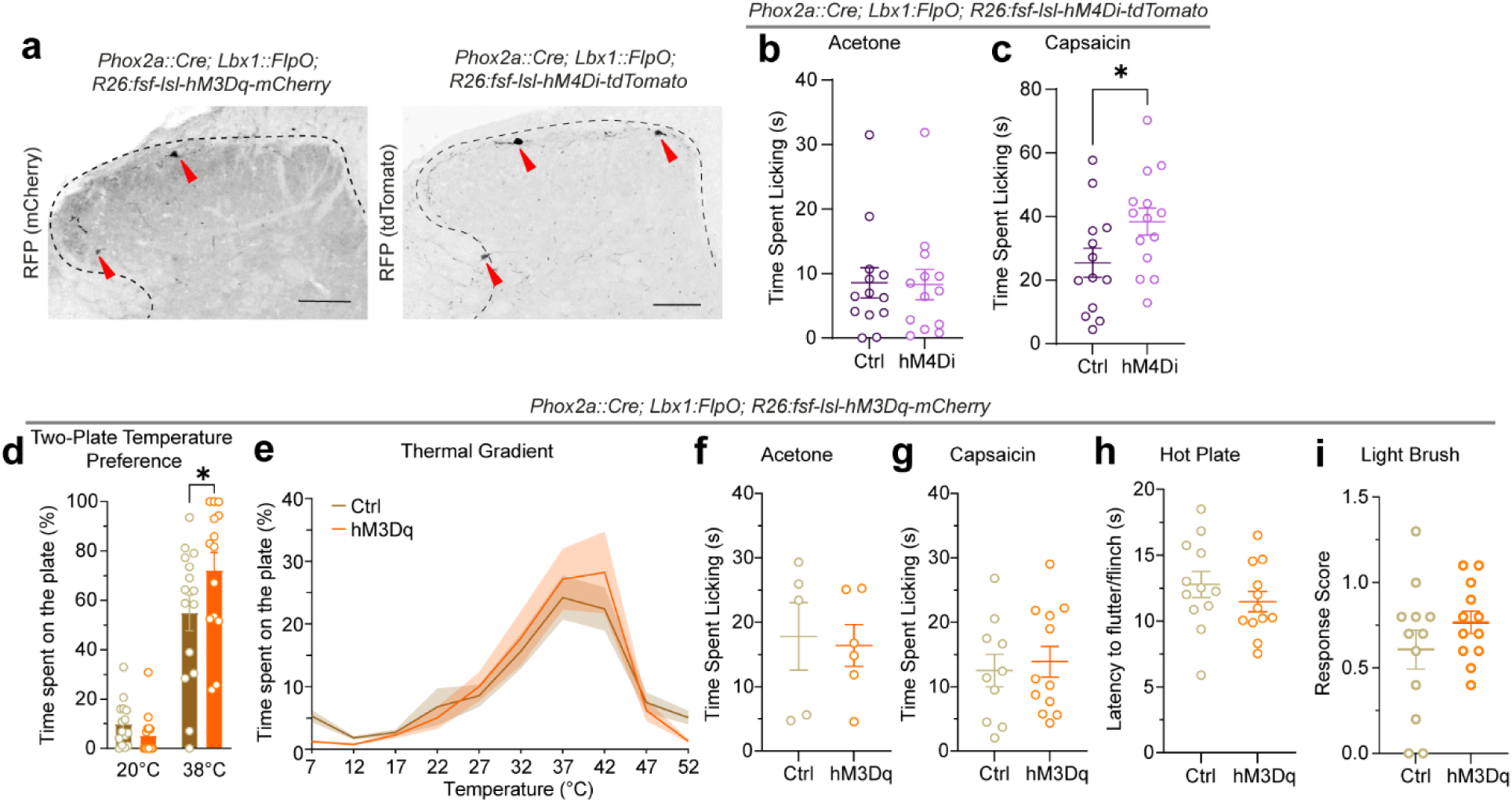
Characterization of chemogenetic models and their behaviours. **a,** Expression of DREADDs (hM3Dq and hM4Di) in the *Phox2a::Cre; Lbx1:FlpO* intersected Phox2a ALT neurons (red arrow). **b, c,** Chemogenetic inhibition did not alter licking response to acetone **(b)** but increased licking following intraplanter injection of capsaicin **(c). d, e,** Chemogenetic activation did not affect behavior in a the two-plate temperature preference at 20°C **(d)** or thermal gradient **(e)** assays. **f-i,** Chemogenetic activation did not alter response to acetone **(f)**, intraplanter injection of capsaicin **(g)**, hot plate **(h)**, or light brush **(i).** Mean ± SEM. * p<0.05, ** p<0.01, *** p<0.001. Mann-Whitney test **(b, c, f-i)**. Two-way ANOVA with Bonferroni’s multiple comparisons **(d, e)**.

**Extended Data Fig. 8.**
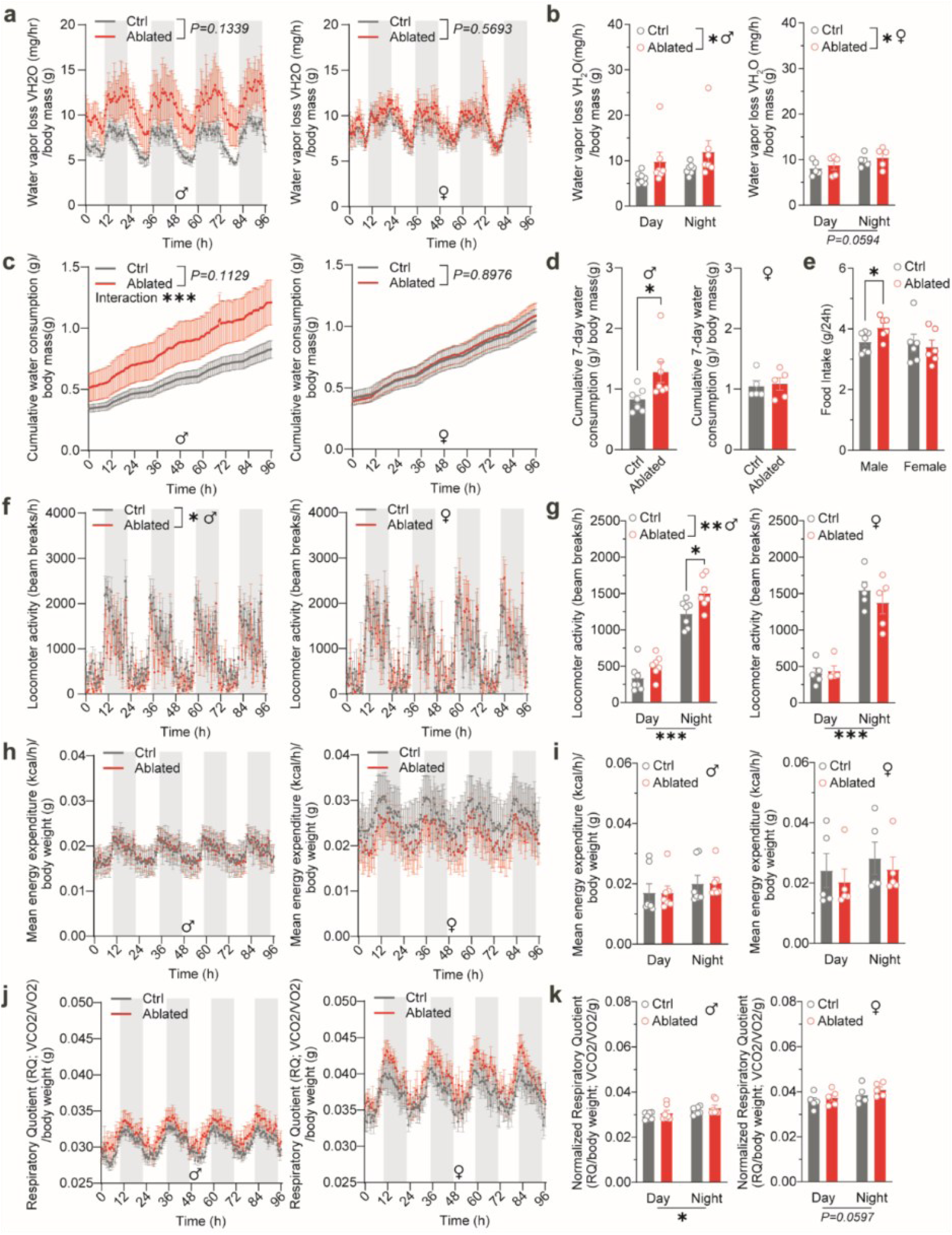
Metabolic parameters of Ablated mcie were largely unaffected. **a, b,** Normalized water vapor loss (VH_2_O) of males and females across a four-day time scale **(a)** and averaged by daytime versus nighttime **(b)**. **c, d,** Normalized cumulative water consumption of males and females across a four-day time scale **(c)** and the total weight of the last measurement **(d)**. **e,** Averaged food intake per 24 hours of males and females. **f, g,** Hourly locomotor activity by number of beam breaks of males and females across a four-day time scale **(f)** and averaged by daytime versus nighttime **(g)**. **h, i,** Normalized energy expenditure of males and females across a four-day time scale **(h)** and averaged by daytime versus nighttime **(i)**. **j, k,** Normalised respiratory quotient (RQ) of males and females across a four-day time scale **(j)** and averaged by daytime versus nighttime **(k)**. Mean ± SEM. * p<0.05, ** p<0.01, *** p<0.001. Two-way ANOVA Repeated Measurement **(a, c, f, h, j)**. Ordinary Two-way ANOVA with Bonferroni’s multiple comparisons **(b, e, g, i, k)**. Mann-Whitney test **(d)**.

**Extended Data Fig. 9.**
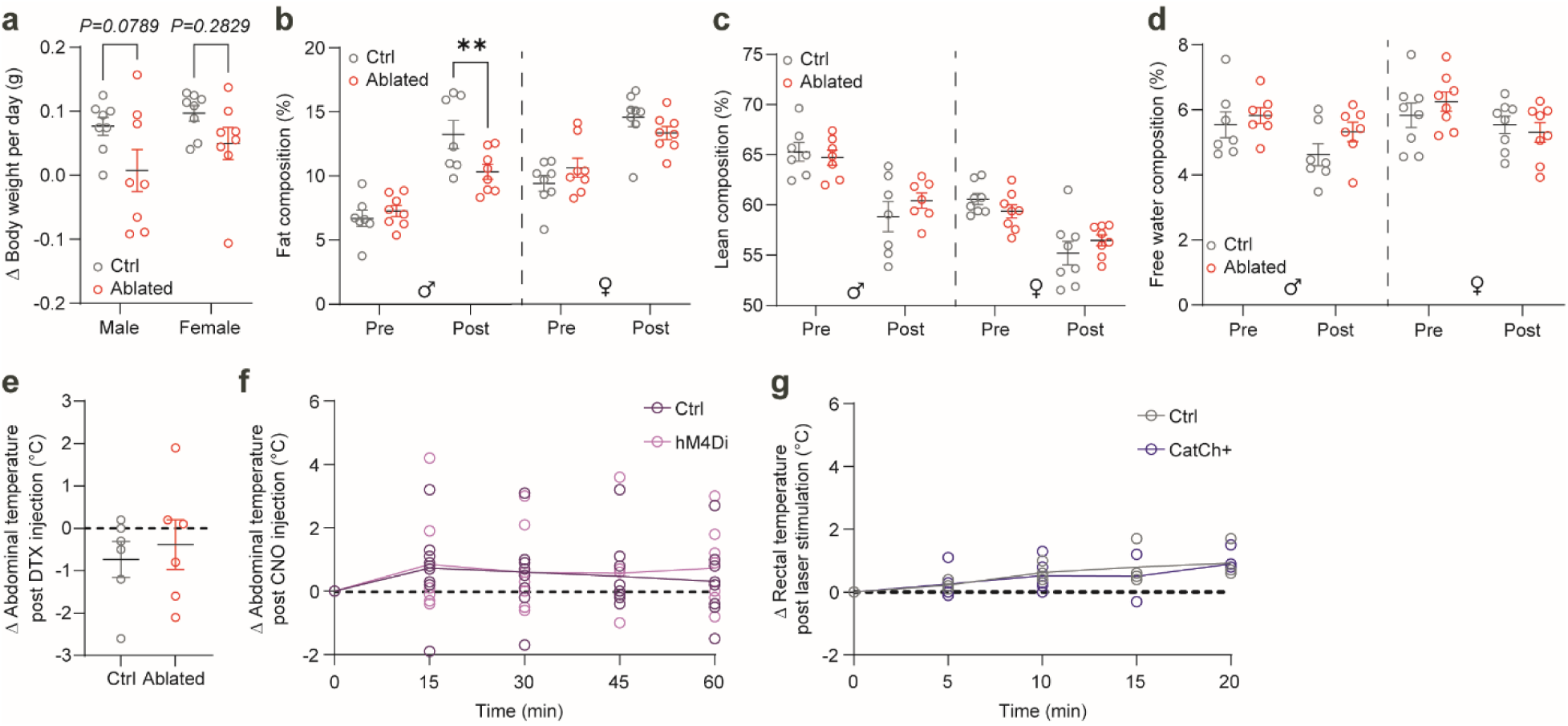
Body composition and core temperature changes in Ablated and chemogenetic model mice. **a,** Averaged body weight changes per day of males and females. **b-d,** Body composition of fat **(b)**, lean **(c)**, and free water **(d)** of male and female mice pre- and 30 days post-injection of DTX. **e, f,** Core body temperatures changes measured by abdominal-implanted chips (UIDTM) 30 days post-injection of DTX **(e)**, 60 minutes post-injection of CNO **(f),** or measured by rectal probe every 5 minutes after optogenetic activation for 20 minutes **(g)**. Mean ± SEM. * p<0.05, ** p<0.01, *** p<0.001. Ordinary Two-way ANOVA with Bonferroni’s multiple comparisons **(a)**, Mann-Whitney test **(e)**, Two-way ANOVA Repeated Measurement with Bonferroni’s multiple comparisons **(b-d, f, g)**.

